# Age-related changes of Peak width Skeletonized Mean Diffusivity (PSMD) across the adult life span: a multi-cohort study

**DOI:** 10.1101/2020.01.07.896951

**Authors:** Grégory Beaudet, Ami Tsuchida, Laurent Petit, Christophe Tzourio, Svenja Caspers, Jan Schreiber, Zdenka Pausova, Yash Patel, Tomas Paus, Reinhold Schmidt, Lukas Pirpamer, Perminder S. Sachdev, Henry Brodathy, Nicole Kochan, Julian Trollor, Wei Wen, Nicola J. Armstrong, Ian J. Deary, Mark E. Bastin, Joanna M. Wardlaw, Susana Munõz Maniega, A. Veronica Witte, Arno Villringer, Marco Duering, Stéphanie Debette, Bernard Mazoyer

**Affiliations:** Institute of Neurodegenerative Diseases (IMN), CNRS, CEA, & University of Bordeaux, Bordeaux, France; Bordeaux Population Health Research Center, Inserm U1219 & University of Bordeaux, Bordeaux, France; Institute of Neuroscience and Medicine (INM-1), Research Centre Juelich, Juelich, Germany; Institute for Anatomy I, Medical Faculty, Heinrich Heine University Dusseldorf, Dusseldorf, Germany; The Hospital for Sick Children and Department of physiology and Nutritional Sciences, University of Toronto, Toronto, ON, Canada; Bloorview Research Institute, Holland Bloorview Kids Rehabilitation Hospital and Departments of Psychology & Psychiatry, University of Toronto, Toronto, ON, Canada; Department of Neurology, Medical University of Graz, Graz, Austria; Centre for Healthy Brain Ageing (CHeBA), School of Psychiatry, UNSW Medicine, University of New South Wales, Sydney, NSW, Australia & Neuropsychiatric Institute Prince of Wales Hospital, Randwick, NSW, Australia; Mathematics and Statistics, Murdoch University, Perth, WA, Australia; Centre for Cognitive Ageing and Cognitive Epidemiology, Department of Psychology, University of Edinburgh, Edinburgh, United Kingdom; Brain Research Imaging Centre, Division of Clinical Neurosciences, MRC Institute for Dementia Research, University of Edinburgh, Edinburgh, United Kingdom; Max Planck Institute for Human Cognitive and Brain Sciences, Leipzig, Germany; Institute for Stroke and Dementia Research (ISD), University Hospital, LMU Munich, Munich, Germany; Department of Neurology, Bordeaux University Hospital, Bordeaux, France

**Keywords:** ageing, white matter, neurodegeneration, MRI, diffusion, PSMD

## Abstract

Parameters of water diffusion in white matter derived from diffusion-weighted imaging (DWI), such as fractional anisotropy (FA), mean, axial, and radial diffusivity (MD, AD and RD), and more recently, peak width of skeletonized mean diffusivity (PSMD), have been proposed as potential markers of normal and pathological brain ageing. However, their relative evolution over the entire adult lifespan in healthy individuals remains partly unknown during early and late adulthood, and particularly for the PSMD index. Here, we gathered and meta-analyzed cross-sectional diffusion tensor imaging (DTI) data from 10 population-based cohort studies in order to establish the time course of white matter water diffusion phenotypes from post-adolescence to late adulthood. DTI data were obtained from a total of 20,005 individuals aged 18.1 to 92.6 years and analyzed with the same pipeline for computing DTI metrics. For each individual MD, AD, RD, and FA mean values were computed over their FA volume skeleton, PSMD being calculated as the 90% peak width of the MD values distribution across the FA skeleton. Mean values of each DTI metric were found to strongly vary across cohorts, most likely due to major differences in DWI acquisition protocols as well as pre-processing and DTI model fitting. However, age effects on each DTI metric were found to be highly consistent across cohorts. RD, MD and AD variations with age exhibited the same U-shape pattern, first slowly decreasing during post-adolescence until the age of 30, 40 and 50, respectively, then progressively increasing until late life. FA showed a reverse profile, initially increasing then continuously decreasing, slowly until the 70’s, then sharply declining thereafter. By contrast, PSMD constantly increased, first slowly until the 60’s, then more sharply. These results demonstrate that, in the general population, age affects PSMD in a manner different from that of other DTI metrics. The constant increase in PSMD throughout the entire adult life, including during post-adolescence, indicates that PSMD could be an early marker of the ageing process.

## 1 Introduction

Parameters of water diffusion in white matter derived from diffusion-weighted imaging (DWI), such as fractional anisotropy (FA), mean, axial, and radial diffusivity (MD, AD and RD) are well-established markers of normal brain maturation(1)(2)(3)(4)(5) and ageing (6)(7)(8)(9)(10)(11)(12) and have been proposed as potential tools for the investigation of various brain disorders (13)(14)(15)(16)(17)(18)(19).

More recently, peak width of skeletonized mean diffusivity (PSMD, (20)), a new phenotype of white matter microstructure that can be derived from DWI, has been proposed as an imaging biomarker of small vessel disease (SVD, (20)(21)) and a correlate of cognitive impairment, particularly processing speed (20)(22)(21). So far, our knowledge of the PSMD distribution in healthy individuals has been limited to these three previously mentioned studies that all included people aged over 50 years. In addition, none of these studies addressed the issue of changes in PSMD across lifespan, which is critical for establishing whether PSMD could be used as an imaging marker of brain aging as well as an early predictor of age-related disorders or to serve as a tool to monitor outcomes in clinical trials. Here, we gathered and meta-analyzed cross-sectional diffusion tensor imaging (DTI) data from 10 population-based cohort studies in order to establish the time course, from post-adolescence to late adulthood, of the PSMD distribution and compare it with that of more commonly used white matter water diffusion phenotypes in white matter.

## 2 Materials and Methods

### 2.1 Participants

Ten independent datasets coming from cross-sectional cohort studies were gathered in the present study, namely MRi-Share, BIL&GIN, SYS, LIFE-Adult, 1000 BRAINS, UKBiobank, ASPSF, OATS, LBC1936, MAS; (see acronym definition in Table 1 caption). All but three (LIFE-Adult, 1000BRAINS, and UKBiobank) were part of the BRIDGET Consortium (BRain Imaging, cognition, Dementia and next generation GEnomics: a Transdisciplinary approach to search for risk and protective factors of neurodegenerative disease), supported by EU-JPND (European Union Joint Programme – Neurodegenerative Disease Research). The 10 datasets included a total of 20,005 individuals (age range: 18.1 to 92.6 years, 10,807 women and 9,198 men). Tables 1 and 2 detail sample size and age distribution for the 10 cohorts that were all of cross.

**Table 1.** Basic statistics for the 10 contributing datasets. MRi-Share: Magnetic Resonance imaging subcohort of Internet-based Students HeAlth Research Enterprise; BIL&GIN: Brain Imaging of Lateralization study at Groupe d’Imagerie Neurofonctionnelle; SYS cohort: Saguenay Youth Study; ASPSF: Austrian Stroke Prevention Study Family; OATS: Older Australian Twin Study; LBC 1936: Lothian Birth Cohort; MAS: Memory and Ageing Study. TIV: total intracranial volume estimated with Freesurfer 5.3 version except “*” with FreeSurfer 6.0 version. ^†^ data subsample size < 30 (not included in the age category statistical analysis)

| Dataset |  | MRi-Share | BIL&GIN | SYS | LIFE | 1000 BRAINS | UKBiobank | ASPSF | OATS | LBC 1936 | MAS |
| --- | --- | --- | --- | --- | --- | --- | --- | --- | --- | --- | --- |
| Sample size | 20,005 | 1,824 | 410 | 512 | 1,906 | 1,209 | 12,397 | 277 | 386 | 672 | 412 |
|  | (% females) | 72% | 51% | 54% | 51% | 44% | 53% | 60% | 64% | 47% | 53% |
| Age (years) | mean (SD) | 22.1 (2.3) | 26.9 (8.0) | 49.5 (5.0) | 58.1 (15.1) | 60.8 (13.4) | 62.6 (7.4) | 65.0 (11.1) | 71.1 (5.1) | 72.7 (0.7) | 80.3 (4.6) |
|  | range | [18.1, 34.9] | [18.1, 57.2] | [36.4, 65.4] | [19.0, 82.0] | [18.5, 85.4] | [45.1, 80.3] | [38.5, 85.6] | [65.4, 89.5] | [71.0, 74.2] | [72.7, 92.6] |
| TIV (cm<sup>3</sup>) | females | 1515 (104)* | 1323 (93) | 1348 (164) | 1637 (115) | 1401 (104) | 1466 (203)* | 1396 (110) | 1369 (133) | 1353 (100) | 1267 (141) |
|  | mean (SD) males | 1703 (118)* | 1503 (127) | 1622 (131) | 1831 (131) | 1570 (118) | 1576 (147)* | 1572 (118) | 1574 (138) | 1537 (112) | 1460 (163) |
| N per age category | 18 to 28 (2,200) | 1,778 | 285 | 0 | 101 | 36 | 0 | 0 | 0 | 0 | 0 |
|  | 28 to 38 (340) | 46 | 84 | 3<sup>†</sup> | 142 | 65 | 0 | 0 | 0 | 0 | 0 |
|  | 38 to 48 (707) | 0 | 24<sup>†</sup> | 176 | 250 | 93 | 126<sup>††</sup> | 38 | 0 | 0 | 0 |
|  | 48 to 58 (4,287) | 0 | 17<sup>†</sup> | 311 | 218 | 176 | 3,525 | 40 | 0 | 0 | 0 |
|  | 58 to 68 (6,535) | 0 | 3<sup>†</sup> | 22<sup>†</sup> | 511 | 431 | 5,375 | 52<sup>††</sup> | 144<sup>††</sup> | 0 | 0 |
|  | 68 to 78 (5,515) | 0 | 0 | 0 | 656 | 358 | 3,351 | 129 | 195 | 672 | 154 |
|  | 78 to 98 (421) | 0 | 0 | 0 | 28<sup>†</sup> | 50 | 20<sup>†</sup> | 18<sup>†</sup> | 47 | 0 | 258 |

**Table 2.** Diffusion-weighted imaging acquisition the 10 contributing datasets. (see Table1 legend for the meaning of dataset abbreviated names and supplementary material section for a more detailed description of these datasets). *: multi-shell acquisition.

| Dataset | MRi-Share | BIL&GIN | SYS | LIFE | 1000 BRAINS | UKBiobank | ASPSF | OATS | LBC 1936 | MAS |
| --- | --- | --- | --- | --- | --- | --- | --- | --- | --- | --- |
| MRI scanner | SIEMENS PRISMA | PHILIPS ACHIEVA | SIEMENS | SIEMENS VERIO | SIEMENS TIM TRIO | SIEMENS SKYRA | SIEMENS TIM TRIO | PHILIPS (x2) SIEMENS(x2) | GE Signa Horizon HDxt | PHILIPS ACHIEVA Quasar Dual |
| B0 strength | 3T | 3T | 1.5T | 3T | 3T | 3T | 3T | 1.5T & 3T | 1.5T | 3T |
| b-gradient (s/mm<sup>2</sup>) | 1000* | 1000 | 1000 | 1000 | 1000 | 1000* | 1000 | 1000 | 1000 | 700 or 1000 |
| Nr of directions | 32 | 2 x 21 | 64 | 60 | 30 | 50 | 6 or 12 | 32 | 64 | 6 or 32 |
| Voxel size (mm<sup>3</sup>) | 1.7x1.7x1.8 | 2x2x2 | 2.3x2.3x3 | 1.7x1.7x1.7 | 2x2x2 | 2x2x2 | 1.8x1.8x2.5 | 2.5x2.5x2.5 | 2x2x2 | 2.5x2.5x2.5 |
| TR (ms) | 3540 (multiband x3) | 8500 | 8000 | 13,800 | 7800 | 3600 (multiband x3) | 4900 or 6700 | 7800 or 8600 | 16500 | 7800 |
| TE (ms) | 75 | 81 | 102 | 100 | 83 | 92 | 81 or 95 | 68 or 96 | 98 | 68 |
| Acquisition duration | 9'45 | 7'45 | 10' | 16'08 | 4'30 | 9'45 | 4'51 or 6'10 | 4'40 or 4'56 | 19'30 | 4'40 |

### 2.2 Diffusion-weighted image acquisition and preprocessing

Tables 3 and 4 summarize the main characteristics of the DWI acquisition and preprocessing for the 10 cohorts. Overall, there was considerable variability between studies regarding almost all acquisition parameters, including scanner manufacturer, field strength, gradient strength, diffusion pulse sequence, resolution and number of directions. For the present work, it was not possible to access raw DWI data at different sites in order to harmonize processing from the initial DICOM data. For this reason, DWI datasets were pre-processed with procedures specific to each site, including exclusion of data upon visual detection of major artifacts due to eddy current distortions or head motion. AD, RD, MD and FA maps were computed by fitting the DTI model parameters in each voxel from these preprocessed DWI volumes. Additional details on DWI preprocessing and DTI parameter map computation for each dataset are provided in the Supplementary Material section.

### 2.3 Derivation of DTI metrics

Various metrics were derived from the DTI data using a script developed by Baykara et al. (http://www.psmd-marker.com, (20)). This original script was designed to extract PSMD, an index of the dispersion of MD values across the white matter skeleton. Briefly, the computation included two steps: 1-WM skeletonizing using the FA map, and 2-analyzing the voxel value distribution histogram in the MD volume masked by the WM-FA skeleton. The FA volume of each individual was skeletonized using the FSL-TBSS software, part of the FMRIB Software Library (FSL) (23)(24), using the FMRIB 1mm FA template and applying a 0.2 threshold on FA maps. Then the MD volume of the same individual was masked, keeping only voxels within the FA skeleton. Furthermore, in order to reduce contamination of the skeleton by CSF voxels, the FA-masked MD volumes were further masked by both a standard FA skeleton with a threshold of 0.3 on FA values and a custom mask (provided with the PSMD software tool) designed so as to exclude regions adjacent to the ventricles, such as the fornix. Finally, PSMD was computed as the difference between the 95^th^ and 5^th^ percentiles of the so-masked MD volume voxel value distribution. Here, we extended this script in order to obtain, in addition to PSMD values, estimates of the mean values of axial, radial, and mean diffusivity (AD, RD, MD, respectively) as well as of fractional anisotropy (FA) over the same customized skeleton. All 10 cohorts were processed separately with this customized script and the results sent to the Bordeaux site where they were combined for further statistical analysis.

### 2.4 Statistical analyses

#### 2.4.1 Age category definition

Due to previously reported non-linear effects of age on DTI metrics (1)(3)(8), we divided each cohort sample into subsamples of 10-years age range starting at 18 years of age, the last subsample (i.e. [78 to 98]) including all subjects aged over 78 years, as there was only a small number of individuals aged over 88 years. Table 1 and Figure 1 detail the contribution of each cohort to each age category. Because we planned running analyses at the age category by cohort by sex level, we discarded subsamples of small sizes, namely containing less than 30 individuals.

**Figure 1.**
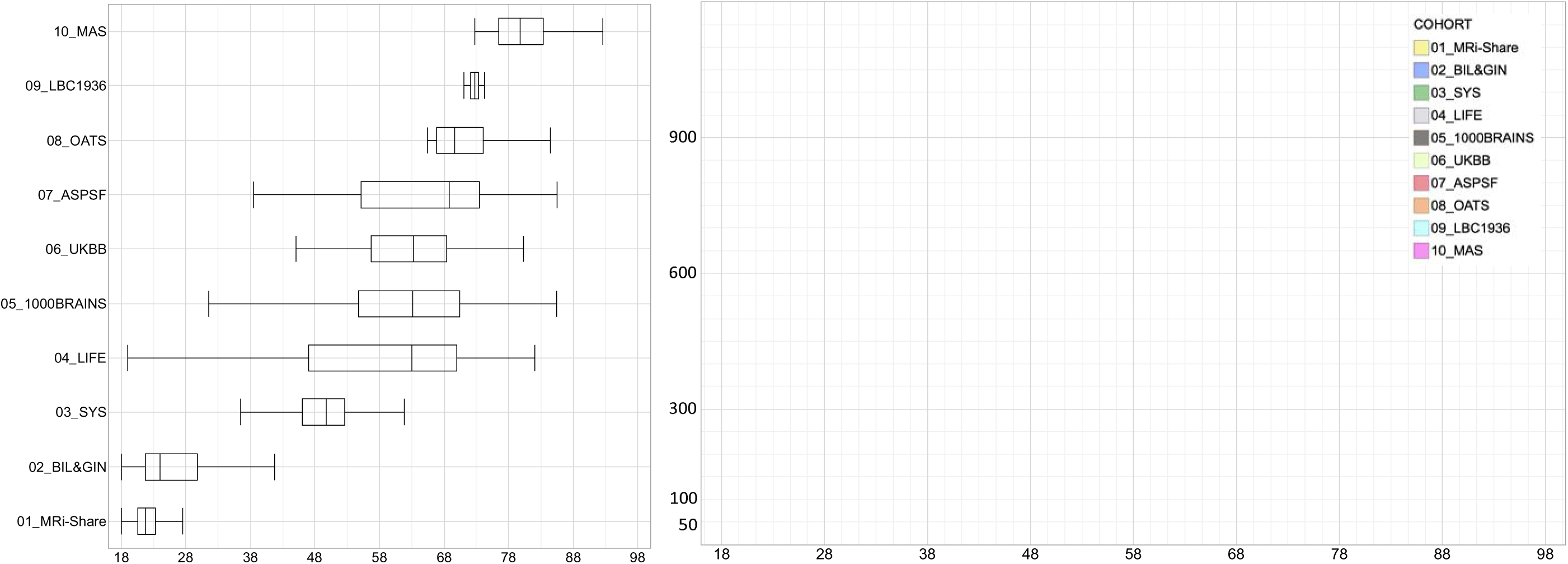
Box-plots and histogram distribution of the age of the participants of the 10 contributing datasets.

#### 2.4.2 Assessing age-related changes of PSMD and other DTI metrics

For each of the five DTI metrics and each age category, we performed an analysis of variance including “Age” as the main effect, and “Sex”, “TIV” (total intracranial volume), and “Cohort” as confounding factors. The Cohort effect was included in order to account for apparent large differences in DTI metric average values across cohorts contributing to the same age category dataset (see Figure 2). In order to document differences of age effects between cohorts contributing to the same age category, we also performed an analysis of variance for each age category and each cohort, including “Age”, “Sex” and “TIV” as effects. Moreover, we analyzed the effects of age on DTI metrics in men and women separately.

**Figure 2.**
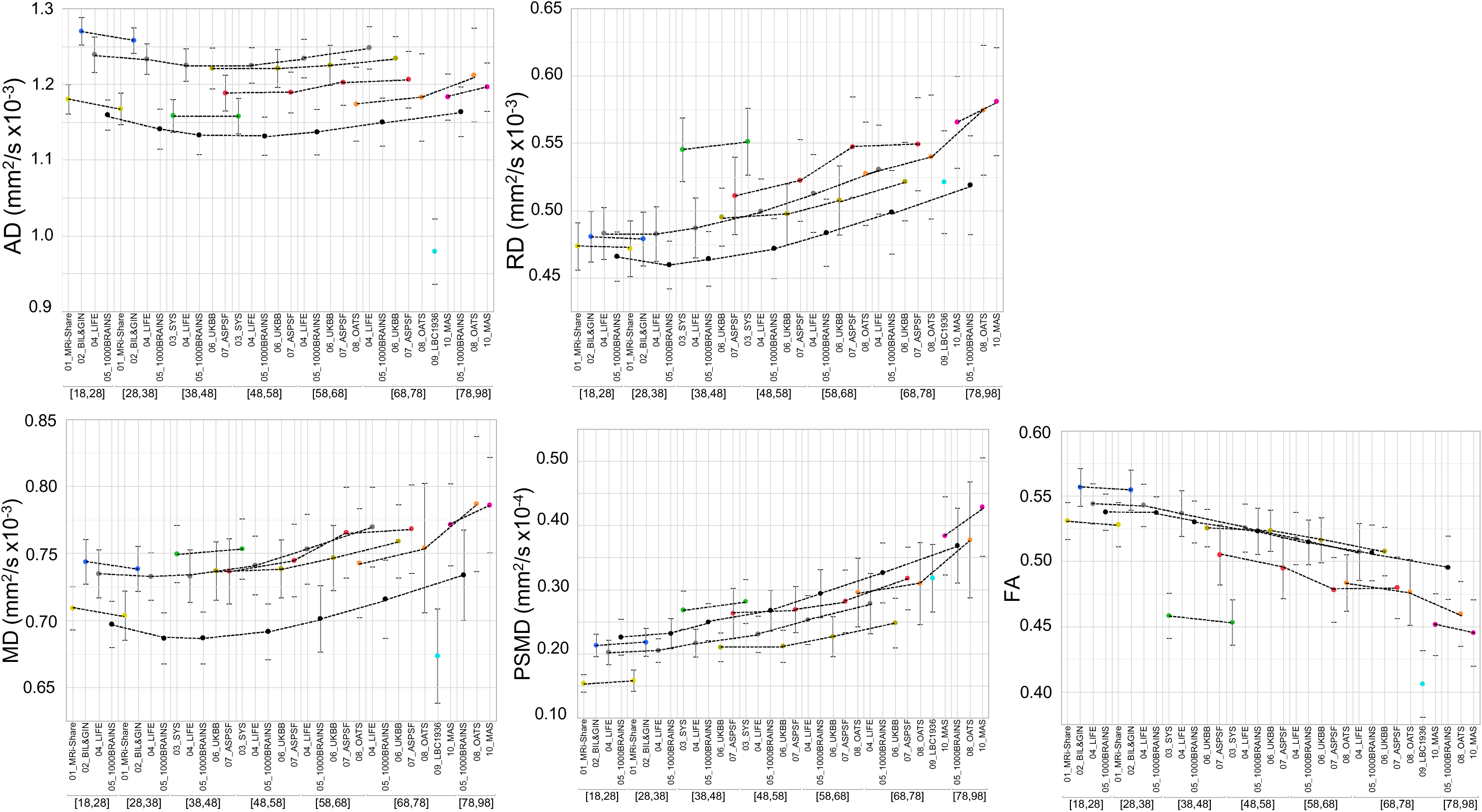
Mean and standard deviation (bars) of the five DTI metrics for each age subcategory and each dataset. The dotted lines connect values for the different age categories of the same dataset.

#### 2.4.3 Assessing the effects of Sex and TIV on PSMD and other DTI metrics

For each of the five DTI metrics and each age category, we also performed an analysis of variance including “Sex” and “TIV” as main factors and “Cohort” as confounding factors.

All statistical analyses were performed using the JMP Pro Software (version 14.3.0, SAS Institute Inc.).

## 3 Results

### 3.1 Descriptive statistics

Tables 3a to 3e provide basic statistics across age categories and cohorts for PSMD and the four other DTI metrics, while Figure 2 illustrates their respective profiles across the adult lifespan. From both Table 3 and Figure 2 it is apparent that there is considerable variability in all five DTI parameter values, and that within a given age category the variability across cohorts is larger than the variability between individuals of the same cohort (see the extreme case of AD and FA values for the LBC1936 study performed at 1.5T, for example).

**Table 3a.** Basic statistics of axial diffusivity (AD, in mm^2^sec^-1^ x 10^-4^) average across a white matter skeleton for each age category and each contributing dataset. Values are mean (S.D. and range [min, max] across the age category sample). See Table1 legend for the meaning of dataset abbreviated names

| Age range | MRi-Share | BIL&GIN | SYS | LIFE | 1000 BRAINS | UKBiobank | ASPSF | OATS | LBC 1936 | MAS |
| --- | --- | --- | --- | --- | --- | --- | --- | --- | --- | --- |
| 18 to 28 | 11.80 (0.19)<br>[10.87, 12.45] | 12.70 (0.18)<br>[12.26, 13.18] |  | 12.39 (0.23)<br>[11.70, 13.10] | 11.60 (0.20)<br>[11.09, 11.96] |  |  |  |  |  |
| 28 to 38 | 11.68 (0.21)<br>[11.10, 12.12] | 12.58 (0.17)<br>[12.14, 13.04] |  | 12.33 (0.20)<br>[11.80, 12.90] | 11.41 (0.27)<br>[10.71, 11.95] |  |  |  |  |  |
| 38 to 48 |  | 12.45 (0.21)<br>[12.07, 12.88] | 11.58 (0.22)<br>[11.03, 12.29] | 12.25 (0.21)<br>[11.60, 12.80] | 11.33 (0.25)<br>[10.52, 11.89] | 12.21 (0.27)<br>[10.76, 12.85] | 11.88 (0.24)<br>[11.50, 12.46] |  |  |  |
| 48 to 58 |  |  | 11.58 (0.23)<br>[11.01, 12.38] | 12.25 (0.24)<br>[11.40, 13.10] | 11.31 (0.25)<br>[10.59, 12.02] | 12.21 (0.25)<br>[10.18, 13.03] | 11.89 (0.27)<br>[11.42, 12.68] |  |  |  |
| 58 to 68 |  |  | 11.67 (0.24)<br>[11.24, 12.21] | 12.34 (0.26)<br>[11.60, 13.20] | 11.37 (0.30)<br>[10.30, 12.32] | 12.25 (0.26)<br>[10.52, 13.24] | 12.03 (0.30)<br>[11.41, 12.76] | 11.74 (0.49)<br>[10.61, 12.80] |  |  |
| 68 to 78 |  |  |  | 12.48 (0.28)<br>[11.40, 13.40] | 11.50 (0.32)<br>[10.36, 12.58] | 12.34 (0.29)<br>[10.70, 13.53] | 12.06 (0.37)<br>[9.69, 12.88] | 11.83 (0.58)<br>[10.69, 13.18] | 9.79 (0.43)<br>[8.41, 10.94] | 11.83 (0.31)<br>[11.06, 12.56] |
| 78 to 98 |  |  |  | 12.49 (0.30)<br>[11.90, 13.20] | 11.64 (0.33)<br>[11.07, 12.47] | 12.44 (0.27)<br>[11.66, 12.83] |  | 12.12 (0.62)<br>[10.80, 13.49] |  | 11.96 (0.32)<br>[10.99, 13.05] |

**Table 3b.** Basic statistics of radial diffusivity (RD, in mm^2^sec^-1^ x 10^-4^) average across a white matter skeleton for each age category and each contributing dataset. Values are mean (S.D. and range [min, max] across the age category sample. (see Table1 legend for the meaning of dataset abbreviated names)

| Age range | MRi-Share | BIL&GIN | SYS | LIFE | 1000 BRAINS | UKBiobank | ASPSF | OATS | LBC 1936 | MAS |
| --- | --- | --- | --- | --- | --- | --- | --- | --- | --- | --- |
| **18 to 28** | 4.74 (0.18)<br>[4.15, 5.53] | 4.81 (0.19)<br>[4.26, 5.44] |  | 4.83 (0.19)<br>[4.33, 5.29] | 4.66 (0.18)<br>[4.26, 5.19] |  |  |  |  |  |
| **28 to 38** | 4.72 (0.21)<br>[4.35, 5.10] | 4.79 (0.20)<br>[4.45, 5.56] |  | 4.83 (0.20)<br>[4.42, 5.36] | 4.60 (0.18)<br>[4.20, 5.17] |  |  |  |  |  |
| **38 to 48** |  | 4.83 (0.22)<br>[4.49, 5.27] | 5.45 (0.24)<br>[4.88, 6.36] | 4.87 (0.22)<br>[4.35, 5.59] | 4.64 (0.20)<br>[4.28, 5.27] | 4.95 (0.22)<br>[4.19, 5.64] | 5.11 (0.29)<br>[4.54, 5.79] |  |  |  |
| **48 to 58** |  |  | 5.51 (0.25)<br>[4.93, 6.54] | 4.99 (0.24)<br>[4.47, 6.01] | 4.72 (0.22)<br>[4.21, 5.38] | 4.97 (0.23)<br>[4.17, 6.22] | 5.22 (0.30)<br>[4.57, 6.05] |  |  |  |
| **58 to 68** |  |  | 5.58 (0.31)<br>[5.02, 6.27] | 5.13 (0.29)<br>[4.29, 6.36] | 4.84 (0.25)<br>[4.21, 5.57] | 5.08 (0.26)<br>[4.34, 6.40] | 5.47 (0.37)<br>[4.73, 6.59] | 5.27 (0.38)<br>[4.36, 6.05] |  |  |
| **68 to 78** |  |  |  | 5.31 (0.33)<br>[4.48, 6.73] | 4.99 (0.31)<br>[4.16, 6.29] | 5.21 (0.29)<br>[4.29, 7.10] | 5.49 (0.35)<br>[4.85, 6.69] | 5.40 (0.46)<br>[4.44, 6.77] | 5.21 (0.38)<br>[4.28, 8.87] | 5.66 (0.34)<br>[5.05, 6.78] |
| **78 to 98** |  |  |  | 5.37 (0.33)<br>[4.87, 6.31] | 5.19 (0.37)<br>[4.69, 5.99] | 5.36 (0.20)<br>[5.03, 5.69] |  | 5.75 (0.48)<br>[4.80, 7.08] |  | 5.81 (0.40)<br>[4.88, 7.17] |

**Table 3c.** Basic statistics of mean diffusivity (MD, in mm^2^sec^-1^ x 10^-4^) average across a white matter skeleton for each age category and each contributing dataset. Values are mean (S.D. and range [min, max] across the age category sample. (see Table1 legend for the meaning of dataset abbreviated names)

| Age range | MRi-Share | BIL&GIN | SYS | LIFE | 1000 BRAINS | UKBiobank | ASPSF | OATS | LBC 1936 | MAS |
| --- | --- | --- | --- | --- | --- | --- | --- | --- | --- | --- |
| 18 to 28 | 7.09 (0.16)<br>[6.54, 7.76] | 7.44 (0.17)<br>[7.02, 7.98] |  | 7.35 (0.18)<br>[6.89, 7.75] | 6.97 (0.17)<br>[6.54, 7.34] |  |  |  |  |  |
| 28 to 38 | 7.04 (0.18)<br>[6.64, 7.38] | 7.39 (0.17)<br>[7.08, 7.97] |  | 7.33 (0.18)<br>[6.92, 7.78] | 6.87 (0.19)<br>[6.43, 7.39] |  |  |  |  |  |
| 38 to 48 |  | 7.37 (0.20)<br>[7.05, 7.80] | 7.50 (0.21)<br>[6.98, 8.17] | 7.33 (0.20)<br>[6.81, 7.84] | 6.87 (0.19)<br>[6.42, 7.39] | 7.37 (0.22)<br>[6.38, 8.05] | 7.37 (0.25)<br>[6.97, 8.01] |  |  |  |
| 48 to 58 |  |  | 7.53 (0.23)<br>[6.97, 8.45] | 7.41 (0.22)<br>[6.89, 8.31] | 6.92 (0.21)<br>[6.35, 7.58] | 7.39 (0.22)<br>[6.17, 8.42] | 7.45 (0.27)<br>[6.91, 8.26] |  |  |  |
| 58 to 68 |  |  | 7.61 (0.27)<br>[7.09, 8.20] | 7.53 (0.26)<br>[6.82, 8.64] | 7.01 (0.25)<br>[6.35, 7.78] | 7.47 (0.24)<br>[6.41, 8.67] | 7.66 (0.34)<br>[6.98, 8.58] | 7.43 (0.41)<br>[6.55, 8.18] |  |  |
| 68 to 78 |  |  |  | 7.70 (0.30)<br>[6.93, 8.92] | 7.16 (0.29)<br>[6.30, 8.39] | 7.59 (0.27)<br>[6.43, 9.23] | 7.68 (0.33)<br>[6.51, 8.74] | 7.54 (0.48)<br>[6.57, 8.91] | 6.74 (0.35)<br>[5.79, 7.86] | 7.72 (0.31)<br>[7.14, 8.71] |
| 78 to 98 |  |  |  | 7.74 (0.31)<br>[7.23, 8.61] | 7.34 (0.34)<br>[6.84, 8.15] | 7.72 (0.20)<br>[7.34, 8.01] |  | 7.87 (0.51)<br>[6.92, 9.21] |  | 7.86 (0.36)<br>[6.96, 9.06] |

**Table 3d.** Basic statistics of peak width skeletonized mean diffusivity (PSMD, in mm^2^sec^-1^ x 10^-4^) for each age category and each contributing dataset. Values are mean (S.D. and range [min, max] across the age category sample. (see Table1 legend for the meaning of dataset abbreviated names.

| Age range | MRi-Share | BIL&GIN | SYS | LIFE | 1000 BRAINS | UKBiobank | ASPSF | OATS | LBC 1936 | MAS |
| --- | --- | --- | --- | --- | --- | --- | --- | --- | --- | --- |
| 18 to 28 | 1.54 (0.14)<br>[1.20, 2.13] | 2.13 (0.17)<br>[1.75, 2.78] |  | 2.02 (0.19)<br>[1.68, 2.53] | 2.26 (0.28)<br>[1.86, 3.47] |  |  |  |  |  |
| 28 to 38 | 1.58 (0.17)<br>[1.33, 2.01] | 2.18 (0.22)<br>[1.75, 2.85] |  | 2.05 (0.19)<br>[1.65, 2.63] | 2.32 (0.24)<br>[1.75, 3.00] |  |  |  |  |  |
| 38 to 48 |  | 2.24 (0.18)<br>[1.89, 2.58] | 2.68 (0.30)<br>[2.07, 4.44] | 2.17 (0.22)<br>[1.64, 3.00] | 2.49 (0.29)<br>[1.94, 3.46] | 2.10 (0.22)<br>[1.60, 3.11] | 2.63 (0.39)<br>[1.95, 3.53] |  |  |  |
| 48 to 58 |  |  | 2.82 (0.34)<br>[1.99, 5.12] | 2.31 (0.28)<br>[1.70, 3.63] | 2.67 (0.32)<br>[2.03, 4.07] | 2.12 (0.25)<br>[1.49, 4.58] | 2.69 (0.36)<br>[2.18, 3.70] |  |  |  |
| 58 to 68 |  |  | 2.92 (0.30)<br>[2.51, 3.57] | 2.53 (0.38)<br>[1.83, 4.85] | 2.94 (0.38)<br>[2.14, 5.03] | 2.27 (0.31)<br>[1.56, 5.71] | 2.82 (0.49)<br>[2.15, 4.54] | 2.96 (0.54)<br>[2.11, 4.90] |  |  |
| 68 to 78 |  |  |  | 2.78 (0.47)<br>[1.88, 5.26] | 3.26 (0.47)<br>[2.23, 5.53] | 2.48 (0.39)<br>[1.61, 6.19] | 3.17 (0.49)<br>[2.15, 4.60] | 3.10 (0.64)<br>[2.17, 6.66] | 3.18 (0.53)<br>[2.09, 5.72] | 3.84 (0.61)<br>[2.56, 6.02] |
| 78 to 98 |  |  |  | 2.96 (0.45)<br>[2.15, 3.94] | 3.68 (0.58)<br>[2.83, 5.32] | 2.74 (0.33)<br>[2.10, 3.45] |  | 3.77 (0.90)<br>[2.47, 6.51] |  | 4.29 (0.77)<br>[2.95, 7.96] |

**Table 3e.** Basic statistics of fractional anisotropy (FA, unitless) average across individual FA skeleton for each age range category and each contributing dataset. Values are mean (S.D. and range [min, max] across the age category sample. (see Table1 legend for the meaning of dataset abbreviated names)

| Age range | MRi-Share | BIL&GIN | SYS | LIFE | 1000 BRAINS | UKBiobank | ASPSF | OATS | LBC 1936 | MAS |
| --- | --- | --- | --- | --- | --- | --- | --- | --- | --- | --- |
| 18 to 28 | 0.53 (0.01)<br>[0.47, 0.58] | 0.56 (0.01)<br>[0.51, 0.60] |  | 0.54 (0.02)<br>[0.49, 0.58] | 0.54 (0.01)<br>[0.49, 0.56] |  |  |  |  |  |
| 28 to 38 | 0.53 (0.02)<br>[0.49, 0.56] | 0.55 (0.02)<br>[0.50, 0.58] |  | 0.54 (0.02)<br>[0.50, 0.58] | 0.54 (0.01)<br>[0.49, 0.57] |  |  |  |  |  |
| 38 to 48 |  | 0.55 (0.02)<br>[0.52, 0.57] | 0.46 (0.02)<br>[0.39, 0.50] | 0.54 (0.02)<br>[0.48, 0.58] | 0.53 (0.02)<br>[0.48, 0.56] | 0.53 (0.01)<br>[0.49, 0.56] | 0.51 (0.02)<br>[0.46, 0.55] |  |  |  |
| 48 to 58 |  |  | 0.45 (0.02)<br>[0.40, 0.50] | 0.53 (0.02)<br>[0.46, 0.57] | 0.52 (0.02)<br>[0.47, 0.58] | 0.52 (0.02)<br>[0.44, 0.57] | 0.49 (0.02)<br>[0.42, 0.54] |  |  |  |
| 58 to 68 |  |  | 0.45 (0.02)<br>[0.41, 0.49] | 0.52 (0.02)<br>[0.44, 0.58] | 0.51 (0.02)<br>[0.46, 0.57] | 0.52 (0.02)<br>[0.44, 0.57] | 0.48 (0.02)<br>[0.41, 0.53] | 0.48 (0.02)<br>[0.43, 0.55] |  |  |
| 68 to 78 |  |  |  | 0.51 (0.02)<br>[0.40, 0.56] | 0.51 (0.02)<br>[0.44, 0.56] | 0.51 (0.02)<br>[0.37, 0.57] | 0.48 (0.02)<br>[0.41, 0.53] | 0.48 (0.03)<br>[0.38, 0.54] | 0.41 (0.03)<br>[0.31, 0.47] | 0.45 (0.02)<br>[0.36, 0.49] |
| 78 to 98 |  |  |  | 0.50 (0.02)<br>[0.45, 0.54] | 0.50 (0.02)<br>[0.44, 0.53] | 0.50 (0.01)<br>[0.48, 0.52] |  | 0.46 (0.02)<br>[0.40, 0.51] |  | 0.45 (0.03)<br>[0.34, 0.51] |

Figure 3 compares the inter-individual variability of PSMD (in terms of its coefficient of variation, CV in %) within each cohort to those of the other DTI metrics, again for each age category and each cohort, revealing that PSMD CV’s is in the order of 10 to 15% (with values as high as 20% for later ages) while those for AD, RD, MD and FA are in the order of 2 to 5%. Note also that the CV’s of all DTI metrics increase with age, more for PSMD than for the other parameters.

**Figure 3.**
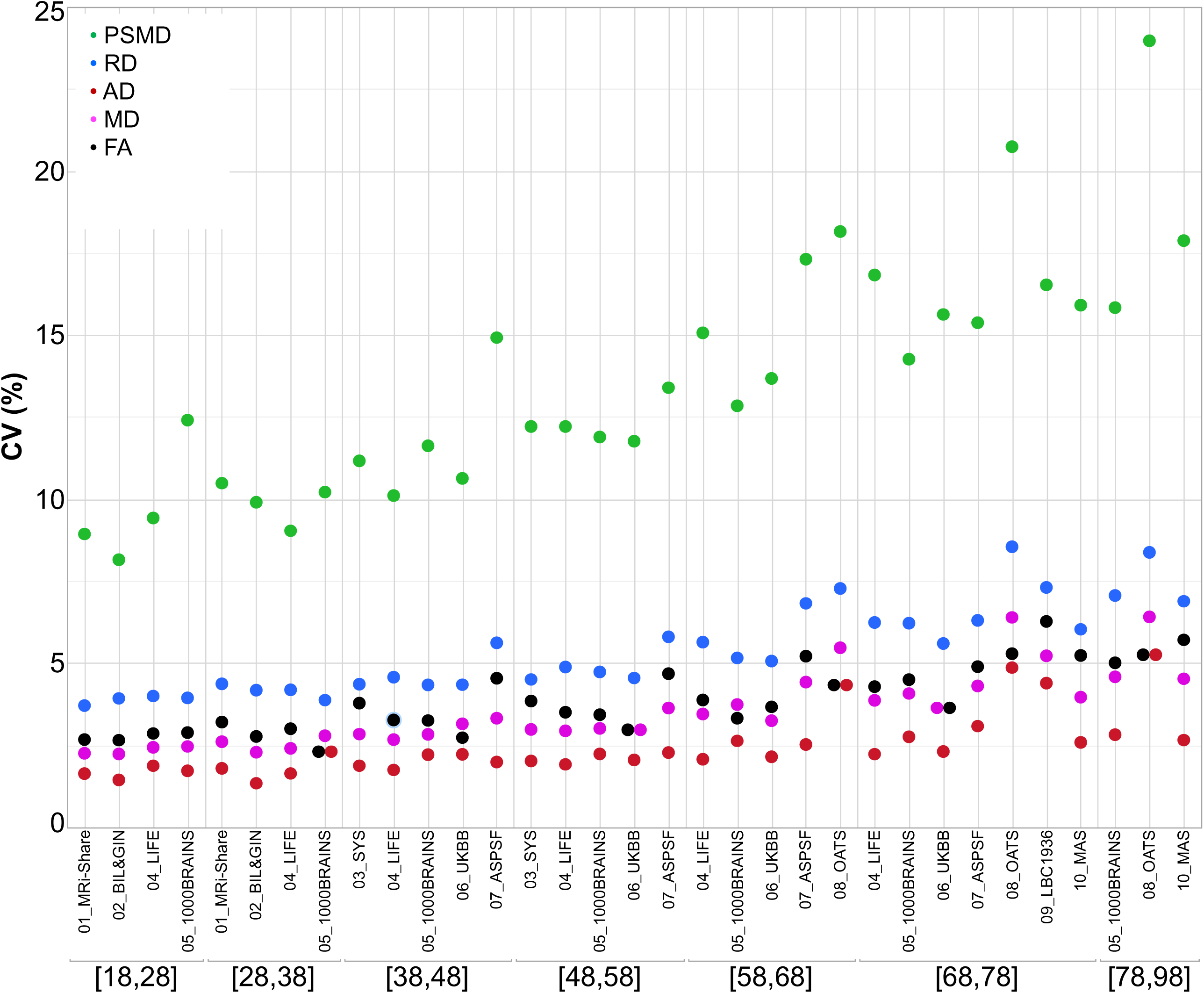
Coefficients of variation of the five DTI metrics for each age subcategory and each dataset.

### 3.2 Effects of age on PSMD and other DTI metrics

The evolution of PSMD across the adult life is different from that of the other metrics (Figure 2). Indeed, PSMD seems to increase monotonically with age whereas AD, RD, and MD exhibit a similar J-shape profile, initially slightly decreasing during post-adolescence before later increasing during adulthood. As for FA, it shows a reverse profile to that of AD, RD and MD, with an initial small increase followed by a later decrease.

This apparent specific lifespan profile of PSMD was confirmed by the quantitative estimates of the effects of age on PSMD and other DTI metrics provided by the between-cohort ANOVA (see Table 4) and their profiles of evolution across age categories as shown in Figure 4. Estimates of the age effect on PSMD were indeed positive for all age categories and significant for all but the [28 to 38] age period. This increase in PSMD accelerated during late life periods, its value being multiplied by a factor of 3 between the [58 to 68] and the [78 to 98] periods. By contrast, AD, RD, and MD age variation profiles were characterized by an initial small but significant decrease (negative age effect), followed by a stable period (non-significant age effect) and a final significant increase (positive age effect). What distinguished AD, RD and MD profiles was their respective timings, the initial decrease in RD and MD being significant only for the [18 to 28] subsample, while it extended over the [38 to 48] period for AD. Note that since the stable period covered two periods of 10 years for all three metrics, the increase in AD was delayed by 10 years compared with RD and MD. At the same time, FA exhibited the reverse contrast consisting of an initial increase during the [18 to 28] period followed by a stable period ([28 to 38]) before an accelerated decrease during the rest of the adult life course. The age effects on DTI metrics were not significantly different between men and women as revealed by the separate sex-specific analyses of age effects.

**Table 4.** ANOVA effect of age (estimate (standard error) and significance p-value) on five DTI metrics (AD, RD, MD, PSMD, and FA evaluated across individual FA skeletons). Values are in mm^2^s^-1^year^-1^ x10^-6^ for AD, RD and MD, in mm^2^s^-1^year^-1^ x10^-7^ for PSMD, and in year^-1^ x10^-3^ for FA.

| Age range (years) | AD |  | RD |  | MD |  | PSMD |  | FA |  |
| --- | --- | --- | --- | --- | --- | --- | --- | --- | --- | --- |
|  | estimate | p | estimate | p | estimate | p | estimate | p | estimate | p |
| 18 to 28 | -2.01 (0.20) | < 0.0001 | -1.06 (0.18) | < 0.0001 | -1.38 (0.17) | < 0.0001 | 0.41 (0.15) | 0.0058 | 0.30 (0.15) | 0.041 |
| 28 to 38 | -1.32 (0.43) | 0.0023 | -0.22 (0.41) | 0.59 | -0.58 (0.37) | 0.12 | 0.60 (0.42) | 0.15 | -0.27 (0.32) | 0.40 |
| 38 to 48 | -0.68 (0.37) | 0.065 | 0.40 (0.37) | 0.27 | 0.047 (0.33) | 0.89 | 1.84 (0.41) | < 0.0001 | -0.54 (0.27) | 0.046 |
| 48 to 58 | -0.11 (0.13) | 0.41 | 0.61 (0.12) | < 0.0001 | 0.37 (0.12) | 0.0019 | 1.11 (0.14) | < 0.0001 | -0.56 (0.09) | < 0.0001 |
| 58 to 68 | 0.95 (0.12) | < 0.0001 | 1.46 (0.11) | < 0.0001 | 1.29 (0.11) | < 0.0001 | 2.18 (0.14) | < 0.0001 | -0.91 (0.08) | < 0.0001 |
| 68 to 78 | 1.75 (0.19) | < 0.0001 | 1.94 (0.18) | < 0.0001 | 1.87 (0.17) | < 0.0001 | 3.67 (0.24) | < 0.0001 | -0.97 (0.12) | < 0.0001 |
| 78 to 98 | 2.30 (0.60) | 0.0002 | 4.01 (0.64) | < 0.0001 | 3.44 (0.59) | < 0.0001 | 6.99 (0.12) | < 0.0001 | -2.36 (0.39) | < 0.0001 |

**Figure 4.**
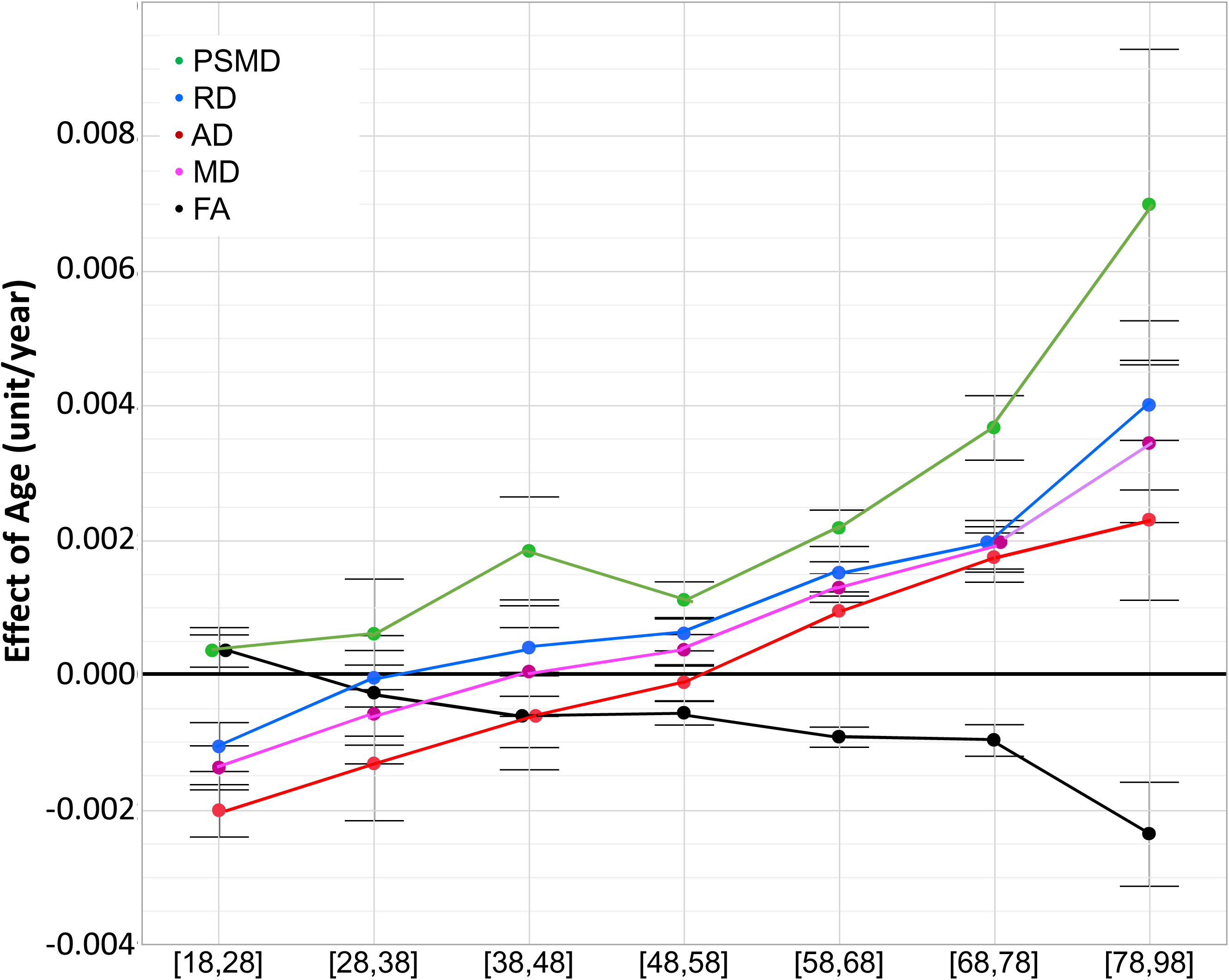
ANOVA estimates and 95% confidence intervals of the effects of age on the five DT metrics for each age subcategory. Units are mm^2^/s/year x10^-3^ for AD, RD and MD, mm^2^/s/year x10^-4^ for PSMD, /year x10^-3^ for FA.

### 3.3 Effects of Sex and TIV on DTI

Amplitude of sex effects on DTI metric average values were found to be quite variable across the various cohorts for the different age categories (see Figure 5). When pooling all datasets, we found that sex had significant effects on all DTI parameters except on FA. Women had higher mean AD and MD values than men, who conversely had higher PSMD values than women (see Table 5). As for TIV, we found that it had positive effects on all DTI parameters (see Table 5), including PSMD, these effects being very significant (p < 10^-4^) in all cases but RD (p=0.53).

**Figure 5.**
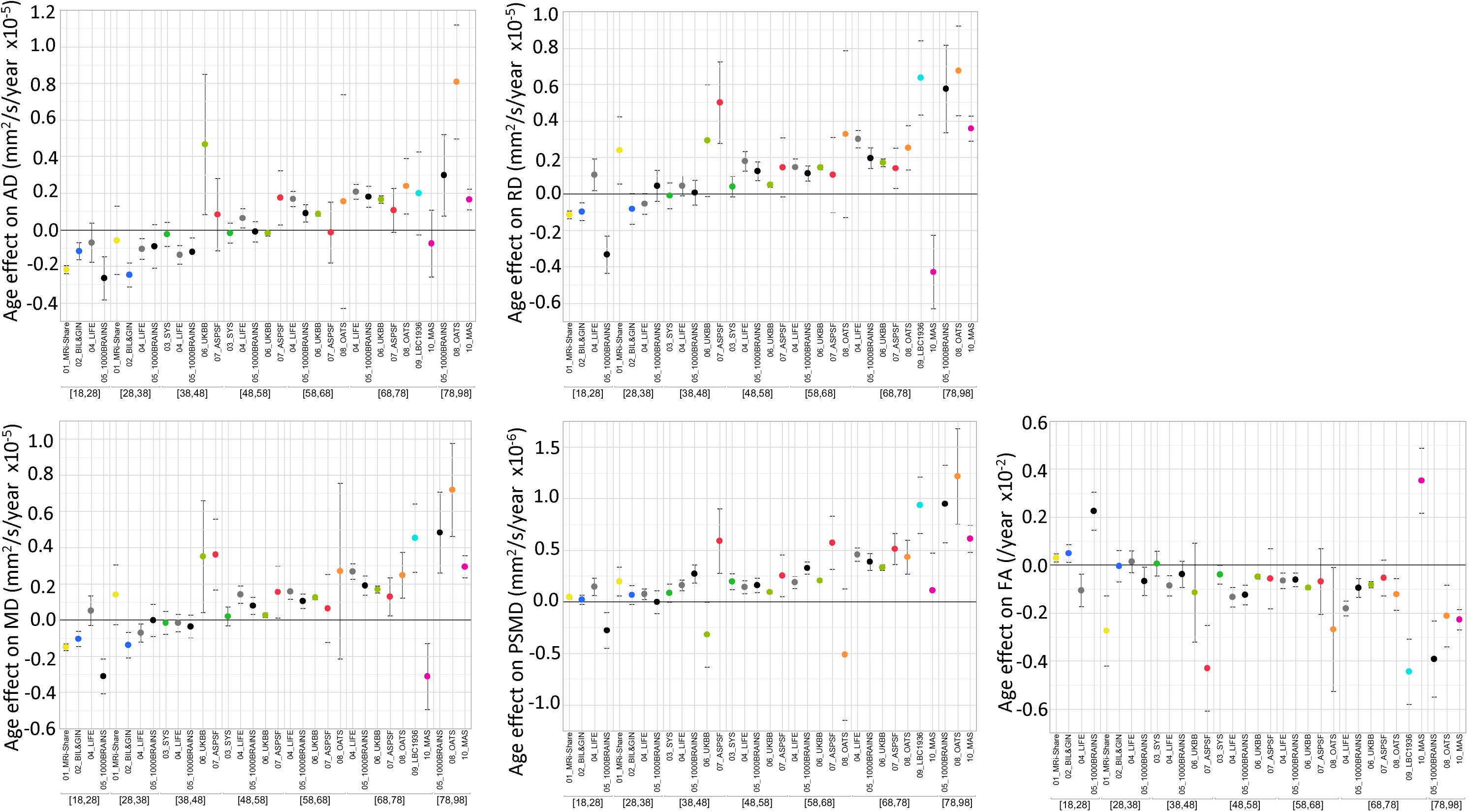
ANOVA estimates and 95% confidence intervals of the effects of age on the five DT metrics for each age subcategory and each dataset.

**Table 5.** ANOVA effect of sex and TIV (estimate (standard error) and significance p-value) on five DTI metrics (AD, RD, MD, PSMD, and FA evaluated across individual FA skeletons). TIV: total intracranial volume. Values for the effects of sex are in mm^2^s^-1^ x10^-6^ for AD, RD, MD, and PSMD and in x10^-3^ for FA, and for the effects of TIV are in mm^2^s^-1^cm^-3^ x10^-6^ for AD, RD MD, and PSMD and in cm^-3^ x10^-5^ for FA.

| Effect | AD |  | RD |  | MD |  | PSMD |  | FA |  |
| --- | --- | --- | --- | --- | --- | --- | --- | --- | --- | --- |
|  | estimate | p | estimate | p | estimate | p | estimate | p | estimate | p |
| Sex (F-M) | 1.13 (0.21) | < 0.0001 | 0.45 (0.21) | 0.032 | 0.68 (0.19) | 0.0005 | -5.34 (0.26) | < 0.0001 | 0.04 (0.14) | 0.74 |
| TIV | 1.41 (0.12) | < 0.0001 | 0.07 (0.12) | 0.53 | 0.52 (0.01) | < 0.0001 | 1.43 (0.15) | < 0.0001 | 0.54 (0.08) | < 0.0001 |

## 4 Discussion

We will first discuss methodological issues and potential limitations in interpreting results of our study. Second, we will discuss what the present study adds to the already existing literature on age effects on classical DTI parameters. In the third part, we will focus on the original findings regarding PSMD distribution and evolution over the adult life.

### 4.1 Methodological issues and potential limitations of the present study

In order to study the effects of age over the entire adult lifespan, we gathered DTI data from 20,005 individuals scanned at 10 different sites with the major objective of maximizing statistical power. By so doing, we were aware that the variability of DTI parameters derived from the entire dataset would be much larger than for a single site study because of between-site differences in data acquisition protocols such as scanner manufacturer, field strength, number and strength of diffusion-encoding gradients, image voxel size, and post-processing of raw DWI data, and so on (23)(24)(25)(26)(27)(28). Accordingly, a key issue was to reduce as much as possible the variability due to sources that were controllable so that we could maximize benefits of the very large sample size.

Concerns with DTI data meta-analysis have been emphasized by several authors (29)(30) and strategies have been proposed for harmonizing DWI data before proceeding to statistical analysis of DTI parameter maps. However, it has also been reported that DTI parameters estimated over the whole white matter or large regions of interest actually exhibit high intra- and inter-scanner reproducibility, making them suitable for multisite studies without extensive harmonization (17)(31)(32). This is the approach that we implemented here as it was not possible to access raw DWI data at different sites in order to harmonize processing from the very earliest stages (see Supplementary Materials). In our study, harmonization was limited to the post-processing of DTI maps pre-computed by each site using the same script for generating individual skeletons and computing individual DTI parameter values across these skeletons. However, our findings indicate that it is possible to draw meaningful conclusions from such minimal harmonization by focusing on the effects of interest (age in our case) rather than the absolute values of the measurements, an approach previously used by others (33).

Multi-site studies do have some advantages as compared with single-site investigations, namely statistical power due to very large sample size, and the ability to recruit a sufficient number of individuals over the entire life or adult lifespan. As emphasized by others, these are important features in the context of clinical studies that are often multisite in nature (17)(31). In fact, there have been several previous reports of multisite studies on DTI parameters in adults but they dealt with issues other than age effects such as methodology (26)(17)(29) and genetic effects (34)(35), for example.

In this study, we focused on the mean DTI metrics over a white matter skeleton, rather than over the entire white-matter compartment, since our primary metric of interest was PSMD, which is defined over the FSL-TBSS white matter skeleton derived from FA data. This approach should in theory minimize any partial volume effects by limiting the measurement over the core of white matter tracts. Indeed, when we compared the mean DTI metrics over the white matter skeleton with those measured over the whole white-matter compartment in two cohorts of the study (MRi-Share and BIL&GIN), mean FA values were higher and diffusivity values lower when averaged over the white matter skeleton than when averaged over the entire white-matter compartment. However, the cross-subject variability measured as the CV of the mean DTI metrics decreased only marginally (a fraction of a percent for CVs ranging from 2% to 4%) when mean values were computed over the white matter skeleton rather than the whole white matter. In contrast, the CV of PSMD decreased markedly (on average from 24% to 8%) when its computation was performed on the white matter skeleton rather than on than the global white-matter mask. This demonstrates the importance of choosing a measure of MD dispersion values over a white matter skeleton for controlling between subject variability.

In the present work, we restricted the analysis to classical DTI metrics as only two of the contributing cohorts had high angular resolution and/or multi-shell acquisition schemes that could be used for estimating advanced white-matter microstructural parameters with more sophisticated models (25). Here, DWI data processing was solely based on the classical DTI model. The DTI model limitations are well-known (24) and it has been shown, for example, that correction for free water has a major impact on classical DTI parameter values (36)(37). However, although investigating advanced white-matter microstructural parameters is highly desirable, it was beyond the scope of our study: it would require additional datasets with multi-shell acquisition, especially for individuals aged 30 to 50 years or over 70 years, in order to supplement existing data (8) on the adult lifespan trajectory of these microstructural parameters.

We included “Sex” and “TIV” as covariates in our statistical analyses. Mixed results have been reported regarding the impact of sex on DTI measures ((38)(39)(40)(12)(41), see review in (3)). Here we also observed mixed results across the different cohorts, although very significant sex effects on all DTI parameters, except FA, were uncovered when combining the entire dataset. Note, however, these sex effects were of very small size (for PSMD, for example, omega^2^=6.7×10^-3^ for the sex effect to be compared with 7.6×10^-2^ for the age effect), which could explain the mixed findings in the literature, and suggests further investigations are required in order to understand their biological origins. TIV effects on DTI parameters are not well established in the literature. In our study, we found that TIV was positively correlated with all DTI parameters except for RD. Similar to sex, TIV effects when significant were very small (for PSMD again, omega^2^=1.5×10^-3^ for the TIV effect). Here again additional investigations are needed to understand the origins of these effects.

Finally, and importantly, it should be stressed that interpretation of the results of the present study should be taken with caution because of the cross-sectional nature of the data that we analyzed. Numerous reports have indeed pointed out the caveats of cross-sectional design for assessing effects of age and demonstrated how such design may lead to spurious findings when compared to those obtained with longitudinal data ((12)(42)(43)). In the present work, there was no attempt to use a single model to describe the variation of DTI parameters with age over the entire adulthood period. Rather, we selected a piecewise linear model to examine/compare age-related changes in 10-year duration consecutive time bins, thereby minimizing the generation bias between cohorts of nearby categories. Understandably, such an approach does not eradicate the intrinsic limits of our cross-sectional study. But it should be reminded that a fully longitudinal design is quasi impossible to implement in the context of lifespan research, since, in practice, measures in an individual can be repeated only a few times and at short duration intervals. As a consequence, such longitudinal studies suffer from some of the limitations of cross-sectional ones. This may explain why the results of the present study are compatible with those a previously published longitudinal study (12) in which individuals aged between 20 and 84 years were observed twice 3 years apart.

### 4.2 Adult life span profiles of variation of classical DTI parameters AD, RD, MD and FA

Effects of age on white-matter microstructure assessed with DTI have been intensively investigated over the past decade from a developmental perspective (4) as well as in a lifespan/ageing framework (1)(11)(40)(3)(12)(11)(5)(8)(36). Briefly, and considering only DTI metrics estimated at the global level, AD, RD and thus MD were reported to follow similar U- or better J-shape age variation patterns, initially decreasing during childhood and adolescence (see (4) for review) then exhibiting an accelerated increase during the adult life (12)(8), while FA followed a reverse profile. Our own findings agree with this body of results during the adult life course. Raw data plots show J-shape profiles for AD, RD and MD, and the reverse profile for FA, as well as acceleration of these changes during late life.

Maximum global FA values and minimum global MD, RD, and AD values have been reported to occur before the age of 40 (1)(40)(11)(5)(12), although large variations were found when considering individual tracts (44)(39)(45)(5). Here, we found extreme values for RD, MD and FA occurring between 28 and 38 years, well in line with these previous findings. In addition, we found that the decrease of AD in the post-adolescence period extended into adulthood by about 10 years more than for RD and MD, thereby uncovering heterochrony of AD and RD variations during adulthood. Such a heterochrony during adulthood was not detected in a previous longitudinal study (12) possibly due to an insufficient sample size and large DTI metric variability between individuals (as can be seen in Fig. 7 of the mentioned report). Note that two recent studies (46)(47) have reported opposite age effects for AD (decrease) and RD (increase) with stable MD during the 18 to 55 year age period; however, as both studies used simple linear modeling due to small sample sizes, no age at extreme value could be observed. Rather, our findings are compatible with the AD-RD variation heterochrony that has been noticed earlier during childhood and adolescence at the individual tract level with stronger decrease for RD than for AD (44)(30). According to these and our findings, the AD decrease / RD increase profile (6) would occur only during mid-adulthood.

### 4.3 PSMD is a diffusion imaging phenotype with a profile of variation across the adult life span that differs from that of other DTI parameters

The distribution of PSMD values observed for the different cohorts and age categories of the present study are consistent with the few comparable data reported in the literature for older participants (no data are available in young adults). For example, Baykara et al. reported in their pioneering article a PSMD median value around 3.0 (in mm^2^s^-1^ x 10^-4^, range [2.5, 4.9]) in a sample of healthy individuals aged 60 to 80 years drawn from the ASPF cohort (see Table 2 of Baykara et al (20)), values that are comparable to those reported in Table 3d of our study in subsamples of other cohorts of similar age category. Similarly, Wei et al (21) recently reported a PSMD average value of 2.4 x 10^-4^ mm^2^s^-1^ in a sample of healthy controls aged around 60 years. In both studies, PSMD CVs were close to 10%, a value again similar to those observed in our own study. That the CV of PSMD is 2 to 3 times larger than the CVs of other DTI metrics could be expected since PSMD is a dispersion rather than a central tendency statistic. Moreover, the larger increase in PSMD CV as age advances (as compared to the other DTI metrics) indicates that this phenotype should be used with caution especially during the late life period. However, it is important to note that the CV of PSMD was found to be quite stable across cohorts with similar age ranges.

The main goal of the present study was to document the profile of PSMD evolution across age bands during adulthood. In this respect, and the proviso that the data we gathered were not longitudinal, our results show that PSMD increases continuously from post-adolescence to late adult life, that this increase is accelerating at later ages, and that this acceleration is larger than for the other DTI metrics. As there are no available data of PSMD in childhood and adolescence, it is not possible to decide whether the lifetime PSMD evolution profile is similar to those of AD, RD, MD, i.e. with a decrease during childhood that reaches the minimum value before adulthood, or if it shows continuous increase throughout the lifespan. Nevertheless, it remains the case that the continuous and accelerating increase of PSMD during adulthood is an indication that it is an adequate and potentially valuable marker of white matter ageing. In particular, it is notable that PSMD increases during early adulthood when the other DTI metrics variations appear to be still undergoing late maturational processes.

The biological mechanisms of the origin of PSMD evolution with age are at present unknown, but one can think of several reasons why PSMD may be more prone to increase with age as compared with the other metrics. First, it is important to remember that PSMD is a measure of MD values dispersion across a skeleton of white matter. As such, it will be directly affected by differences across MD values of the individual tracts. Consequently, regional heterogeneity as well as heterochrony in MD values of the fiber tracts will result in higher PSMD values more than in average MD values. Second, MD itself is a weighted average of AD (1/3) and RD (2/3) values, and thus MD value dispersion will also be affected by heterochrony in AD and RD variations with age. Overall, what possibly makes PSMD an early and sensitive imaging marker of ageing is that it captures multiple sources of heterogeneity in white matter water diffusion parameters. With this regard, it would be interesting to investigate variations in the pattern of MD dispersion at the regional level using tract-based DTI metrics since it is well established that heterochrony is a major feature of the development and aging of the different fiber tracts (see for example (44)(39)(8)(5)). Accordingly, variations of PSMD value provide only a gross and possibly biased estimate of the white matter-microstructure dynamics. We did not implement regional analysis as our study focused on PSMD that is by definition a dispersion statistic over the entire white matter skeleton. Nevertheless, a regional approach would certainly be interesting and feasible since peak width of MD values could be measured on a white-matter skeleton at the tract level in the same manner as it has been done for other DTI metrics (see (30)(46) for example).

## Supporting information

Cohort description

## 5 Acknowledgments

Gregory Beaudet has been supported by an EU-ERC starting grant (SEGWAY, PI S Debette, with funding from the European Union’s Horizon 2020 research and innovation program under grant agreement No 640643). Ami Tsuchida is supported by a grant from the Fondation pour la Recherche Médicale (DIC202161236446). Svenja Caspers was supported by the Initiative and Networking Fund of the Helmholtz Association and the European Union’s Horizon 2020 Research and Innovation Program under Grant Agreement 785907 (Human Brain Project SGA2). The BRIDGET project is supported by the Fondation Leducq (Transatlantic Network of Excellence on the Pathogenesis of SVD of the Brain) and is an EU Joint Program - Neurodegenerative Disease Research (JPND) project. The project is supported through the following funding organizations under the aegis of JPND -www.jpnd.eu: Australia, National Health and Medical Research Council, Austria, Federal Ministry of Science, Research and Economy; Canada, Canadian Institutes of Health Research; France, French National Research Agency; Germany, Federal Ministry of Education and Research; Netherlands, The Netherlands Organization for Health Research and Development; United Kingdom, Medical Research Council. This project has received funding from the European Union’s Horizon 2020 research and innovation program under grant agreement No 643417. This project has also received funding from the European Research Council (ERC) under the European Union’s Horizon 2020 research and innovation program under grant agreement No 640643.

## 6 Author Contributions

Study conception (BM, SD), data collection (GB, CT, SC, ZP, TP, RS, PS, HB, NK, JT, ID, VW, AV, BM), data analysis (GB, LP, SC, JS, YP, LP, PS, WW, NA, MB, SMM, VW, MD, BM), drafting (GB, AT, BM), revising the manuscript (LP, CT, SC, JS, ZP, YP, TP, R, LP, PS, WW, NA, ID, MB, JW, SMM, VW, AV, MD, SD, BM).

## 11 Data Availability Statement

Requests to access these datasets should be directed to:

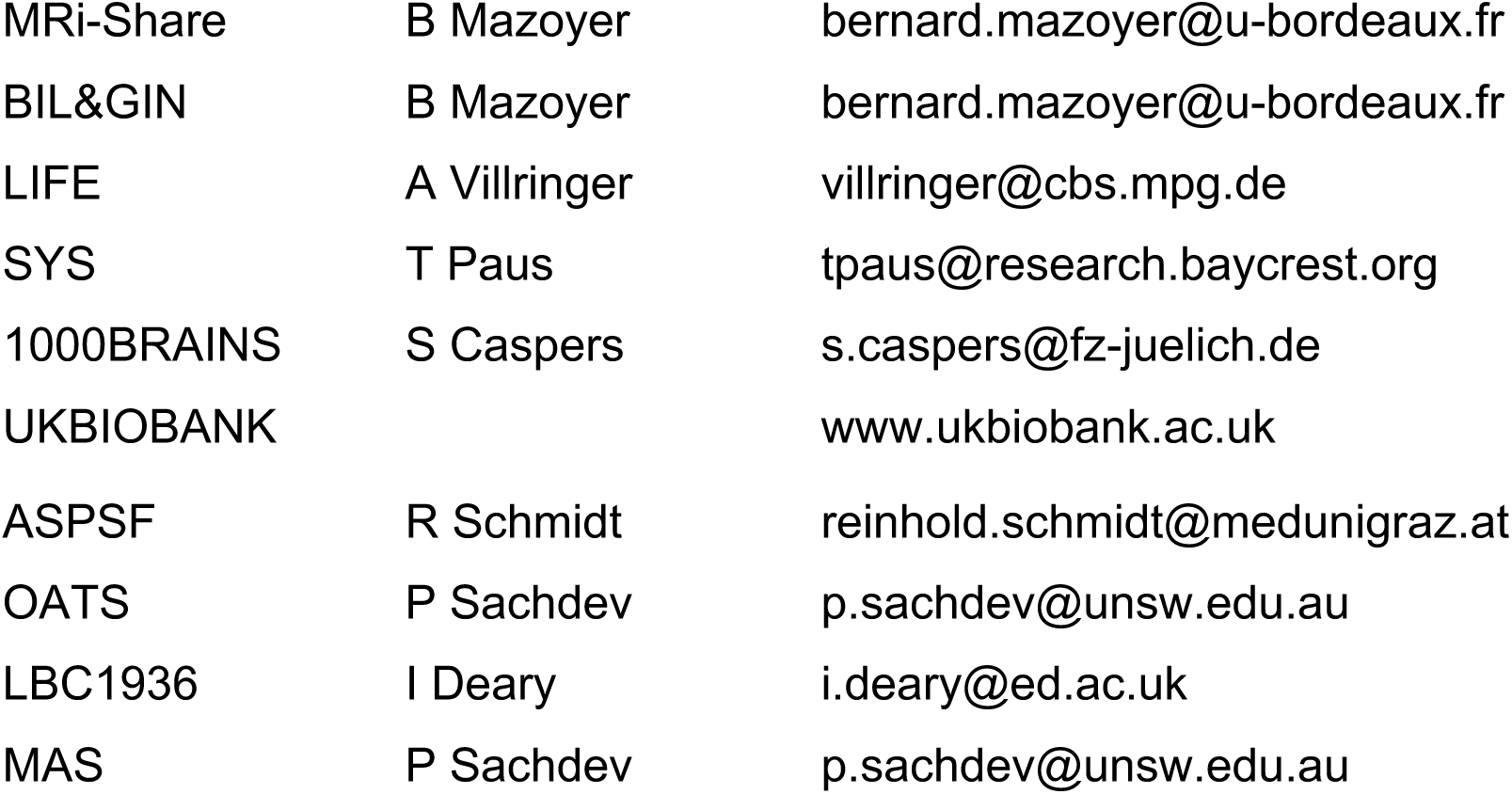

## References

1. Hasan KM, Sankar A, Halphen C, Kramer LA, Brandt ME, Juranek J, Cirino PT, Fletcher JM, Papanicolaou AC, Ewing-Cobbs L. Development and organization of the human brain tissue compartments across the lifespan using diffusion tensor imaging. Neuroreport (2007) 18:1735–1739.

2. Giorgio A, Watkins KE, Chadwick M, James S, Winmill L, Douaud G, De Stefano N, Matthews PM, Smith SM, Johansen-Berg H, et al. Longitudinal changes in grey and white matter during adolescence. NeuroImage (2010) 49:94–103. doi:10.1016/j.neuroimage.2009.08.003

3. Lebel C, Gee M, Camicioli R, Wieler M, Martin W, Beaulieu C. Diffusion tensor imaging of white matter tract evolution over the lifespan. NeuroImage (2012) 60:340–352. doi:10.1016/j.neuroimage.2011.11.094

4. Lebel C, Treit S, Beaulieu C. A review of diffusion MRI of typical white matter development from early childhood to young adulthood. NMR Biomed (2019) 32:e3778. doi:10.1002/nbm.3778

5. Yeatman JD, Wandell BA, Mezer AA. Lifespan maturation and degeneration of human brain white matter. Nat Commun (2014) 5:4932. doi:10.1038/ncomms5932

6. Bennett IJ, Madden DJ, Vaidya CJ, Howard DV, Howard JH. Age-related differences in multiple measures of white matter integrity: A diffusion tensor imaging study of healthy aging. Hum Brain Mapp (2010) 31:378–390. doi:10.1002/hbm.20872

7. Bartzokis G, Lu PH, Heydari P, Couvrette A, Lee GJ, Kalashyan G, Freeman F, Grinstead JW, Villablanca P, Finn JP, et al. Multimodal Magnetic Resonance Imaging Assessment of White Matter Aging Trajectories Over the Lifespan of Healthy Individuals. Biol Psychiatry (2012) 72:1026–1034. doi:10.1016/j.biopsych.2012.07.010

8. Cox SR, Ritchie SJ, Tucker-Drob EM, Liewald DC, Hagenaars SP, Davies G, Wardlaw JM, Gale CR, Bastin ME, Deary IJ. Ageing and brain white matter structure in 3,513 UK Biobank participants. Nat Commun (2016) 7:13629. doi:10.1038/ncomms13629

9. de Lange A-MG, Bråthen ACS, Rohani DA, Fjell AM, Walhovd KB. The temporal dynamics of brain plasticity in aging. Cereb Cort (2018) 28:1857–1865. doi: 10.1093/cercor/bhy003

10. Rathee R, Rallabandi VPS, Roy PK. Age-related differences in white matter integrity in healthy human brain: evidence from structural MRI and diffusion tensor imaging. Magn Reson Insights (2016) 9:9–20. doi:10.4137/MRI.S39666

11. Westlye LT, Walhovd KB, Dale AM, Bjornerud A, Due-Tonnessen P, Engvig A, Grydeland H, Tamnes CK, Ostby Y, Fjell AM. Life-Span Changes of the Human Brain White Matter: Diffusion Tensor Imaging (DTI) and Volumetry. Cereb Cortex (2010) 20:2055–2068. doi:10.1093/cercor/bhp280

12. Sexton CE, Walhovd KB, Storsve AB, Tamnes CK, Westlye LT, Johansen-Berg H, Fjell AM. Accelerated Changes in White Matter Microstructure during Aging: A Longitudinal Diffusion Tensor Imaging Study. J Neurosci (2014) 34:15425–15436. doi:10.1523/JNEUROSCI.0203-14.2014

13. for the Alzheimer’s Disease Neuroimaging Initiative, Chandra A, Dervenoulas G, Politis M. Magnetic resonance imaging in Alzheimer’s disease and mild cognitive impairment. J Neurol (2019) 266:1293–1302. doi:10.1007/s00415-018-9016-3

14. Tae W-S, Ham B-J, Pyun S-B, Kang S-H, Kim B-J. Current clinical applications of diffusion-tensor imaging in neurological disorders. J Clin Neurol (2018) 14:129. doi:10.3988/jcn.2018.14.2.129

15. Shen X, Adams MJ, Ritakari TE, Cox SR, McIntosh AM, Whalley HC. White matter microstructure and its relation to longitudinal measures of depressive symptoms in mid- and late life. Biol Psychiatry (2019) 86:759–768. doi:10.1016/j.biopsych.2019.06.011

16. Suzuki H, Gao H, Bai W, Evangelou E, Glocker B, O’Regan DP, Elliott P, Matthews PM. Abnormal brain white matter microstructure is associated with both pre-hypertension and hypertension. PLOS ONE (2017) 12:e0187600. doi:10.1371/journal.pone.0187600

17. Croall ID, Lohner V, Moynihan B, Khan U, Hassan A, O’Brien JT, Morris RG, Tozer DJ, Cambridge VC, Harkness K, et al. Using DTI to assess white matter microstructure in cerebral small vessel disease (SVD) in multicentre studies. Clin Sci (2017) 131:1361–1373. doi:10.1042/CS20170146

18. Dekkers IA, Jansen PR, Lamb HJ. Obesity, Brain volume, and white matter microstructure at MRI: A Cross-sectional UK Biobank Study. Radiology (2019) 291:763–771. doi:10.1148/radiol.2019181012

19. Zavaliangos-Petropulu A, Nir TM, Thomopoulos SI, Reid RI, Bernstein MA, Borowski B, Jack Jr. CR, Weiner MW, Jahanshad N, Thompson PM. Diffusion MRI Indices and their relation to cognitive impairment in brain aging: the updated multi-protocol approach in ADNI3. Front Neuroinformatics (2019) 13:2. doi:10.3389/fninf.2019.00002

20. Baykara E, Gesierich B, Adam R, Tuladhar AM, Biesbroek JM, Koek HL, Ropele S, Jouvent E, Alzheimer’s Disease Neuroimaging Initiative, Chabriat H, et al. a novel imaging marker for small vessel disease based on skeletonization of white matter tracts and diffusion histograms. Ann Neurol (2016) 80:581–592. doi:10.1002/ana.24758

21. Wei N, Deng Y, Yao L, Jia W, Wang J, Shi Q, Chen H, Pan Y, Yan H, Zhang Y, et al. A Neuroimaging marker based on diffusion tensor imaging and cognitive impairment due to cerebral white matter lesions. Front Neurol (2019) 10:81. doi:10.3389/fneur.2019.00081

22. Deary IJ, Ritchie SJ, Muñoz Maniega S, Cox SR, Valdés Hernández MC, Luciano M, Starr JM, Wardlaw JM, Bastin ME. Brain Peak Width of Skeletonized Mean Diffusivity (PSMD) and cognitive function in later life. Front Psychiatry (2019) 10:524. doi:10.3389/fpsyt.2019.00524

23. Barrio-Arranz G, de Luis-García R, Tristán-Vega A, Martín-Fernández M, Aja-Fernández S. Impact of MR acquisition parameters on DTI scalar indexes: a tractography based approach. PLOS ONE (2015) 10:e0137905. doi:10.1371/journal.pone.0137905

24. Curran KM, Emsell L, Leemans A. “Quantitative DTI Measures,” in Diffusion Tensor Imaging, eds. W. Van Hecke, L. Emsell, S. Sunaert (New York, NY: Springer New York), 65– 87. doi:10.1007/978-1-4939-3118-7_5

25. Alexander DC, Dyrby TB, Nilsson M, Zhang H. Imaging brain microstructure with diffusion MRI: practicality and applications. NMR Biomed (2019) 32:e3841. doi:10.1002/nbm.3841

26. Helmer KG, Chou M-C, Preciado RI, Gimi B, Rollins NK, Song A, Turner J, Mori S. Multi-site study of diffusion metric variability: characterizing the effects of site, vendor, field strength, and echo time using the histogram distance. in Proc SPIE Int Soc Opt Eng (2016), eds. B. Gimi, A. Krol (San Diego, California, United States), 9788. doi:10.1117/12.2217449

27. Qin W, Shui Yu C, Zhang F, Du XY, Jiang H, Xia Yan Y, Cheng Li K. Effects of echo time on diffusion quantification of brain white matter at 1.5T and 3.0T. Magn Reson Med (2009) 61:755–760. doi:10.1002/mrm.21920

28. Hutchinson EB, Avram AV, Irfanoglu MO, Koay CG, Barnett AS, Komlosh ME, Özarslan E, Schwerin SC, Juliano SL, Pierpaoli C. Analysis of the effects of noise, DWI sampling, and value of assumed parameters in diffusion MRI models: Cross-Model Analysis of Noise and DWI Sampling. Magn Reson Med (2017) 78:1767–1780. doi:10.1002/mrm.26575

29. Fortin J-P, Parker D, Tunç B, Watanabe T, Elliott MA, Ruparel K, Roalf DR, Satterthwaite TD, Gur RC, Gur RE, et al. Harmonization of multi-site diffusion tensor imaging data. NeuroImage (2017) 161:149–170. doi:10.1016/j.neuroimage.2017.08.047

30. Pohl KM, Sullivan EV, Rohlfing T, Chu W, Kwon D, Nichols BN, Zhang Y, Brown SA, Tapert SF, Cummins K, et al. Harmonizing DTI measurements across scanners to examine the development of white matter microstructure in 803 adolescents of the NCANDA study. NeuroImage (2016) 130:194–213. doi:10.1016/j.neuroimage.2016.01.061

31. Grech-Sollars M, Hales PW, Miyazaki K, Raschke F, Rodriguez D, Wilson M, Gill SK, Banks T, Saunders DE, Clayden JD, et al. Multi-centre reproducibility of diffusion MRI parameters for clinical sequences in the brain: Multi-centre reproducibility of diffusion MRI using clinical sequences. NMR Biomed (2015) 28:468–485. doi:10.1002/nbm.3269

32. Prohl AK, Scherrer B, Tomas-Fernandez X, Filip-Dhima R, Kapur K, Velasco-Annis C, Clancy S, Carmody E, Dean M, Valle M, et al. Reproducibility of Structural and Diffusion Tensor Imaging in the TACERN Multi-Center Study. Front Integr Neurosci (2019) 13:24. doi:10.3389/fnint.2019.00024

33. Jockwitz C, Mérillat S, Liem F, Oschwald J, Amunts K, Caspers S, Jäncke L. Generalizing age effects on brain structure and cognition: A two-study comparison approach. Hum Brain Mapp (2019) 40:2305–2319. doi:10.1002/hbm.24524

34. Jahanshad N, Kochunov PV, Sprooten E, Mandl RC, Nichols TE, Almasy L, Blangero J, Brouwer RM, Curran JE, de Zubicaray GI, et al. Multi-site genetic analysis of diffusion images and voxelwise heritability analysis: A pilot project of the ENIGMA–DTI working group. NeuroImage (2013) 81:455–469. doi:10.1016/j.neuroimage.2013.04.061

35. Kochunov P, Jahanshad N, Sprooten E, Nichols TE, Mandl RC, Almasy L, Booth T, Brouwer RM, Curran JE, de Zubicaray GI, et al. Multi-site study of additive genetic effects on fractional anisotropy of cerebral white matter: Comparing meta and megaanalytical approaches for data pooling. NeuroImage (2014) 95:136–150. doi:10.1016/j.neuroimage.2014.03.033

36. Chad JA, Pasternak O, Salat DH, Chen JJ. Re-examining age-related differences in white matter microstructure with free-water corrected diffusion tensor imaging. Neurobiol Aging (2018) 71:161–170. doi:10.1016/j.neurobiolaging.2018.07.018

37. Wu Y-C, Field AS, Whalen PJ, Alexander AL. Age- and gender-related changes in the normal human brain using hybrid diffusion imaging (HYDI). NeuroImage (2011) 54:1840– 1853. doi:10.1016/j.neuroimage.2010.09.067

38. Sullivan EV, Adalsteinsson E, Hedehus M, Ju C, Moseley M, Lim KO, Pfefferbaum A. Equivalent disruption of regional white matter microstructure in ageing healthy men and women: Neuroreport (2001) 12:99–104. doi:10.1097/00001756-200101220-00027

39. Hasan KM, Kamali A, Abid H, Kramer LA, Fletcher JM, Ewing-Cobbs L. Quantification of the spatiotemporal microstructural organization of the human brain association, projection and commissural pathways across the lifespan using diffusion tensor tractography. Brain Struct Funct (2010) 214:361–373. doi:10.1007/s00429-009-0238-0

40. Kochunov P, Glahn DC, Lancaster J, Thompson PM, Kochunov V, Rogers B, Fox P, Blangero J, Williamson DE. Fractional anisotropy of cerebral white matter and thickness of cortical gray matter across the lifespan. NeuroImage (2011) 58:41–49. doi:10.1016/j.neuroimage.2011.05.050

41. Hsu J-L, Leemans A, Bai C-H, Lee C-H, Tsai Y-F, Chiu H-C, Chen W-H. Gender differences and age-related white matter changes of the human brain: A diffusion tensor imaging study. NeuroImage (2008) 39:566–577. doi:10.1016/j.neuroimage.2007.09.017

42. Fjell AndersM, Walhovd KB, Westlye LT, Østby Y, Tamnes CK, Jernigan TL, Gamst A, Dale AM. When does brain aging accelerate? Dangers of quadratic fits in cross-sectional studies. NeuroImage (2010) 50:1376–1383. doi:10.1016/j.neuroimage.2010.01.061

43. Pfefferbaum A, Sullivan EV. Cross-sectional versus longitudinal estimates of age-related changes in the adult brain: overlaps and discrepancies. Neurobiol Aging (2015) 36:2563–2567. doi:10.1016/j.neurobiolaging.2015.05.005

44. Lebel C, Walker L, Leemans A, Phillips L, Beaulieu C. Microstructural maturation of the human brain from childhood to adulthood. NeuroImage (2008) 40:1044–1055. doi:10.1016/j.neuroimage.2007.12.053

45. Lebel C, Beaulieu C. Longitudinal Development of Human Brain Wiring Continues from Childhood into Adulthood. J Neurosci (2011) 31:10937–10947. doi:10.1523/JNEUROSCI.5302-10.2011

46. Kodiweera C, Alexander AL, Harezlak J, McAllister TW, Wu Y-C. Age effects and sex differences in human brain white matter of young to middle-aged adults: A DTI, NODDI, and q-space study. NeuroImage (2016) 128:180–192. doi:10.1016/j.neuroimage.2015.12.033

47. Tian L, Ma L. Microstructural Changes of the Human Brain from Early to Mid-Adulthood. Front Hum Neurosci (2017) 11:393. doi:10.3389/fnhum.2017.00393

