## Supplementary material for "Age-related changes of Peak width Skeletonized Mean Diffusivity (PSMD) across the adult life span: a multi-cohort study": Cohort description

frontiers

***Supplementary Material***

***Description of the 10 cohorts***

### 1 MRi-Share

#### 1.1 Participants

1,874 students were recruited from the larger sample of the i-Share epidemiologic study (internet – based Student Health Research Enterprise, N=30,000, www.ishare.fr) designed for investigating students' health, among whom 1,824 had a brain MRI acquisition. This subsample (1,313 females, 511 males) was aged between 18.1 and 34.9 years (mean ± S.D.: 22.1 ± 2.3 years), with a level of schooling superior or equal to 12 years (= French baccalaureate, UK A levels, or US high school diploma).

#### 1.2 Image acquisition

Participants were imaged between 2015 and 2018 on the same Siemens Prisma three Tesla MR scanner (MAGNETOM Prisma, Siemens, Erlangen, Germany). Diffusion-weighted imaging (DWI) data were acquired using a multi-shell multiband x3 sequence with 100 non-collinear diffusion gradient directions (b0/2xd8; b300/d8; b1000/d32; b2000/d60). For the needs of the present study, DTI metrics were computed based on b1000 alone. Imaging parameters were follows: TR = 3,540 ms; TE = 75 ms; voxel size 1.7 x 1.7 x 1.8 $mm^3$; gradients: 80 mT/m – 200 T/m/s; 64 channels head coil. The DWI acquisition time was 9 min 45 s.

#### 1.3 DWI preprocessing

All DWI data were preprocessed with a nipype pipeline combining FMRIB Software Library (FSL) release 5.0.10 (www.fmrib.ox.ac.uk/fsl) (1) and the *dipy* tools. Briefly, data were first corrected for eddy current and top-up distortion using the FSL Eddy tool, with replacement of outlier slices (2)(3), and denoised using *nlmeans* denoising tool (4). We extracted the denoised b0 and b1000 to fit the DTI model and obtained FA, MD, AD, and RD maps in DWI space.

### 2 BIL&GIN

#### 2.1 Participants

The BIL&GIN database has been described in detailed elsewhere (5). Shortly, 453 young healthy volunteers were recruited, the sample being balanced for handedness and sex. Diffusion-weighted imaging (DWI) was available in a subsample of 410 participants (209 females, 201 males) aged between 18.1 and 57.2 years (mean ± S.D.: 26.9 ± 8.0), with a level of schooling superior or equal to 12 years (= French baccalaureate, UK A levels, or US high school diploma).

#### 2.2 Image acquisition

Participants were imaged between 2007 and 2011 on the same Philips Achieva 3 Tesla MRI scanner (Philips Medical Systems, Best, The Netherlands). DWI data were acquired using a single-shot spin-echo echo-planar sequence one b0 map (b = 0 s/mm$^2$) with 21 non-collinear diffusion gradient directions (b = 1000 s/mm$^2$) homogeneously spread over a half sphere, the series of 21 directions being acquired twice by reversing the gradients' polarity. 70 axial slices parallel to the AC-PC plane were acquired from the bottom of the cerebellum to the vertex. Imaging parameters were follows: TR = 8500 ms; TE = 81 ms; angle = 90°; SENSE reduction factor = 2.5; FOV 224 mm; acquisition matrix 112 x 112.2 x 2 x 2 mm$^3$ isotropic voxel. A second series of one b0 map of 42 volumes was acquired leading to a total DWI acquisition time of 15 min 30 s (2 x 7 min 45 s).

##### 2.3 DWI preprocessing

All DWI data were separately processed for each participant by combining FMRIB Software Library (FSL) release 5.0.9 (www.fmrib.ox.ac.uk/fsl) (1) and the command-line tools of Diffusion Toolkit (www.trackvis.org/dtk/). In order to compute the correct gradient table necessary to calculate diffusion tensors for DWI data acquired on Philips MRI scanners, the *DTI_gradient_table_creator* was used. A 12-parameters affine intra-modal image registration was performed with FLIRT to correct for movement and geometrical deformations (6). DWI with polarity inversion (2 × 21 volumes) were geometrically averaged (7). Finally, both series of 42 volumes were arithmetically averaged and processed with Dipy-related tools (8) to produce for each participant voxelwise maps of axial, radial and mean diffusivity, and fractional anisotropy (AD, RD, MD, and FA, respectively).

#### 3 SYS

##### 3.1 Participants

A total of 1,991 participants (1,029 adolescents and 962 parents from 481 families) were recruited from the genetic founder population of the Saguenay Lac St Jean region of Quebec, Canada during Wave 1 of the Saguenay Youth Study (SYS). During Wave 2 (2012-2015), 664 parents participated in the collection of additional data. In this sample, 580 had complete MRI data (T1w, and DTI). After quality control of DTI data, a total of 512 participants were used for this report (234 males, 278 females, mean age = 49.5 years, S.D. = 4.95 years). Details regarding participant requirement have been described previously (9)(10).

##### 3.2 Image acquisition

Participants were imaged on the same Siemens Avanto 1.5 Tesla MR scanner. Diffusion-weighted imaging (DWI) data were acquired using a single-shot, spin-echo, echo planar sequence with twice-refocused balanced diffusion encoding gradients. There were 64 diffusion gradient directions (b = 1000) and 1 non-diffusion weighted direction (b = 0). Imaging parameters were as follows: TR = 8,000 ms; TE = 102 ms; voxel size 2.3 x 2.3 x 3.0 mm$^3$.

##### 3.3 DWI preprocessing

All diffusion weighted images were processed using FMRIB Software Library (FSL version 5.0.9) (6). Briefly, DWI maps underwent 1) correction for eddy current distortion, 2) brain extraction and 3) diffusion tensor model fitting using FSL's: "*eddy_correct*", "*bet*", and "*dtifit*", respectively.

### 4 LIFE-Adult

#### 4.1 Participants

LIFE-Adult was supported by the grants from the European Union, the European Regional Development Fund, the Free State of Saxony within the framework of the excellence initiative, the LIFE–Leipzig Research Center for Civilization Diseases, University of Leipzig (Project Numbers: 713-241202, 14505/2470, 14575/2470, and 100329290). The authors also received grants from the German Research Foundation, contract grant number CRC 1052-A1 to AV and WI 3342/3-1 to AVW. 2,249 adult participants (1169 men, 1080 women; 19-82 years old, mean age 58.8 years, S.D. = 15 years) with usable DWI data were included from the population-based LIFE-Adult study (11).

#### 4.2 Image acquisition

MRI data collection took place in 2011-2014 using a 3T Siemens MRI Magnetom Verio (Siemens Healthcare GmbH, Germany). We used a 32-channel head coil and a double spin-echo encoding sequence ([TR]/[TE]: 13800/100 ms, 72 slices, 60 diffusion directions (b=1000 s/mm$^2$), 7 non-diffusion-weighted volumes (b=0 s/mm$^2$), EPI-factor: 128 (resolution 128x128), FoV: 220 × 220 × 123 mm$^3$, voxel size: (1.7 mm)$^3$; duration: 16 minutes 8 seconds). Parallel imaging was performed with a generalized auto-calibrating partially parallel acquisition (GRAPPA), reconstruction algorithm and an acceleration factor of 2.

#### 4.3 DWI preprocessing

Preprocessing included skull stripping with BET and eddy outlier replacement, part of FSL (12)(2). Next, motion correction and co-registration to ACPC-reoriented T1-weighted image was done with Lipsia (13). Then the diffusion tensor was modeled to estimate FA, RD, AD and MD at each voxel using Lipsia again. Due to insufficient fat suppression that created an artifact in some of the images (sculp ghosting in the parietal/occipital lobe), we provided manually visual ratings of artifact severity (three levels: 'none', 'mild', moderate-severe') for every participant.

### 5 1000BRAINS

#### 5.1 Participants

Subjects were recruited from the 1000BRAINS study, a large population-based cohort study on the variability of structure, function and connectivity in brain aging in relation to environmental and genetic risk factors (14). In total, 1,315 participated in 1000BRAINS within the base acquisition. For the current study, 1,209 subjects with available diffusion-imaging data (see below) which passed visual quality control during multiple steps of the processing pipeline were included. Exclusion of subjects was based on criteria such as insufficient precision of brain mask, bad image registration results or bad image quality. This subsample of 1,209 subjects (537 females, 672 males) was aged between 18.5 and 85.4 years (mean ± S.D.: 60.6 ± 13.4 years).

##### 5.2 Image acquisition

Participants within the 1000BRAINS study were imaged between September 2011 and March 2018 on the same Siemens TIM TRIO 3 Tesla MR scanner (MAGNETOM TRIO, Siemens, Erlangen, Germany) with a 32-channel head coil. Out of the complete scanning protocol of 1000BRAINS (for details see (14), T1-weighted (T1w) structural as well as diffusion-weighted data (DWI) with 30 diffusion directions were used. T1w data were acquired with the following parameters: 176 slices, TR = 2.25 s, TE = 3.03 ms, TI = 900 ms, FoV = 256 × 256 $mm^2$, flip angle = 9°, voxel resolution = 1 $mm^3$. DWI data were acquired using the following parameters: 4 images at b = 0, 30 images at b = 1000 $mm^2$/s, TR=7,800 ms; TE=83 ms; voxel resolution: 2 $mm^3$).

##### 5.3 DWI preprocessing

DWI data were preprocessed with a snakemake (version 3.13.0, 15) pipeline which included the relevant processing tools of FSL the FMRIB Software Library (FSL) release 5.0.9 (www.fmrib.ox.ac.uk/fsl) (1). Briefly, data were first corrected for motion and eddy currents using the FSL Eddy tool (3)(2), and then rigidly aligned to the T1w brain image in MNI space implicitly interpolating the data to 1.25 mm isotropic voxel size. The brain mask was computed from the T1w image using the CAT12 toolbox. FA, MD, AD, and RD maps were then computed using *dtifit* from the FSL toolbox fitting the tensor with weighted least squares.

#### 6 UKBiobank

##### 6.1 Participants

The UKBiobank comprises around 500,000 community-dwelling participants who were initially recruited from across Great Britain between 2006 and 2010, aged 40-69 years. A total of 12,397 datasets were obtained from the UKBiobank (http://www.ukbiobank.ac.uk/; approved application reference number 18359). The subsample (6,558 females, 5,839 males) had DWI, and was aged between 45.1 and 80.3 years (mean ± S.D.: 62.6 ± 7.4 years).

##### 6.2 Image acquisition

Participants were imaged on a single standard Siemens Skyra 3 Tesla scanner. DWI data were acquired using a multiband x3 spin-echo echo-planar sequence with 10 T2-weighted (b = 0 s/mm2) baseline, 50 b = 1,000 s/mm2 and 50 b = 2,000 s/mm2 diffusion-weighted volumes acquired with 100 distinct diffusion-encoding directions. For the needs of the present study, DTI metrics were computed based on b1000 alone. Imaging parameters were follows: TR = 3,600 ms; TE = 92 ms; voxel size 2.0 x 2.0 x 2.0 mm3. The DWI acquisition time was 7 min.

##### 6.3 DWI preprocessing

Images were preprocessed and analyzed with the FMRIB Software Library (FSL) (http://fsl.fmrib.ox.ac.uk/fsl). Gradient distortion correction was applied using tools developed by the Freesurfer and Human Connectome Project groups (https://github.com/Washington-University/Pipelines). The Eddy tool from FSL (http://fsl.fmrib.ox.ac.uk/fsl/fslwiki/EDDY/) was then used to correct the data for head motion and eddy currents. FA, MD AD and RD maps were created by fitting the b= 1000 s/mm. shell into DTIFIT (DTI fitting tool).

### 7 ASPSF

#### 7.1 Participants

The Austrian Stroke Prevention Family Study (ASPS-Fam) is a prospective single-center, community-based study on the cerebral effects of vascular risk factors in a normal aging population of the city of Graz, Austria. ASPS-Fam represents an extension of the Austrian Stroke Prevention Study (ASPS), which was established in 1991(16)(17). As previously described, inclusion criteria were no history of stroke or dementia and a normal neurological examination. Diffusion MRI was applied in 278 subjects (110 males, 168 females), mean age=65.04 years (SD=11.11 years).

#### 7.2 Image acquisition

Magnetic resonance imaging was performed on a 3T whole-body MR system (TimTrio; Siemens Healthcare, Erlangen, Germany) with a 12-channel head coil. Diffusion-weighted-imaging was acquired with 6 (n=135) or 12 (n=143) diffusion-gradient directions and following settings: b-values (0,1000), TR=6700ms, TE=95ms, voxel size = 1.8 x 1.8 x 2.5/3 $mm^3$, gradients: 38 mT/m – 170 mT/m/s.

#### 7.3 DWI preprocessing

DWI preprocessing was performed using *eddy* for motion and distortion correction followed by *dtifit*, both part from the FMRIB Software Library (FSL) release 5.0.10 (www.fmrib.ox.ac.uk/fsl) (1).

### 8 OATS

#### 8.1 Participants

Our study cohort was drawn from Wave 1 of the Older Australian Twins Study (OATS), a study of twins aged 65 years or older living in the three Eastern states of Australia (New South Wales, Victoria and Queensland) primarily recruited from the Australian Twin Registry (ATR). methodology of OATS has previously been described in (18). This study was approved by the ethics committees of the Australian Twin Registry, University of New South Wales, University of Melbourne, Queensland Institute of Medical Research and the South-Eastern Sydney & Illawarra Area Health Service. Informed consent was obtained from all subjects and the methods were carried out in accordance with the relevant guidelines.

#### 8.2 Image acquisition

MRI data were obtained on three 1.5 Tesla scanners and a 3 Tesla scanner owing to the multi-site nature of this study. Siemens Magnetom Avanto and Sonata scanners (Siemens Medical Solutions, Malvern PA, USA) with similar years of manufacture and upgrade were used in centers 2 (Victoria) and 3 (Queensland), respectively. In center 1 (New South Wales), a 1.5 Tesla Philips Gyroscan scanner (Philips Medical Systems, Best, Netherlands) was used initially, followed by a 3 Tesla Philips Achieva Quasar Dual scanner. The acquisition protocols and parameters for the 1.5 Tesla scanners were tested and matched between the centers. For each participant, 32 directional diffusion-weighted imaging (DWI) data were acquired with a single-shot, spin-echo, echo-planar imaging (EPI) sequence: b=700 s/mm2, echo time (TE) = 68 ms, repetition time (TR) = 7767 ms, flip angle = 90$^{o}$, matrix size = 240 × 240, field of view (FOV) = 240 × 240 mm, yielding in-plane resolution of 1 × 1 mm, slice thickness 2.5 mm contiguous axial slices without gap. For the 3 Tesla scanner, b=1000 s/mm2, TR =7115 ms, TE = 70 m, with 32 non-collinear directions used.

##### 8.3 DWI preprocessing

DTI data were pre-processed with the FMRIB's Diffusion Toolbox (FDT) of the FMRIB's Software Library (FSL) version 5.0 (http://www.Fmrib.ox.ac.uk/fsl, Oxford Centre for Functional Magnetic Resonance Imaging of the Brain, Oxford University, UK). First, each subject's raw DTI data were corrected for head movement and eddy current distortions by linearly registering each diffusion-weighted volume to the reference T-weighted non-diffusion b0 image, using the FMRIB's linear image registration (FLIRT). A binary brain mask was created to remove the non-brain tissue using the Brain Extraction Tool (BET) in FSL. Then, diffusion tensor model was fitted to each imaging voxel of the pre-processed DTI data to derive Fractional anisotropy (FA) and mean diffusivity (MD) maps.

The resulting FA images were fed into TBSS, which is also a part of FSL, to carry out the voxel-wise statistical analysis. In brief, the FA maps of all participants were spatially transformed to a 1×1×1 mm MNI (Montreal Neurological Institute) standard space by the nonlinear registration method FNIRT. The nonlinearly registered FA images were further averaged to generate a mean FA image. A non-maximum suppression algorithm was applied afterwards to search the image voxels with highest FA value along the direction perpendicular to the local tract surface to create a mean FA skeleton. We then produced the FA, MD, AD, RD and PSMD maps using the common script designed for the present work.

#### 9 LBC1936

##### 9.1 Participants

The Lothian Birth Cohort 1936 (LBC1936) comprises a sample of 1,091 people who were born in 1936 and were recruited between 2004 and 2007 at about 70 years of age (Wave 1, (19)). All lived independently in the community, were generally healthy, and travelled to a clinical research facility for assessment. Most of them had taken part in the Scottish Mental Survey 1947 at age 11 years. At Wave 1, they undertook extensive cognitive, medical, biomarker, psycho-social, and other assessments. The assessments were repeated at Wave 2, about 3 years later at mean age 73 years, with the addition of a detailed structural brain MRI scan. Of the 886 who returned at Wave 2, over 700 agreed to undertake a brain scan (20). All variables employed here were collected at Wave 2. Ethical approval for the LBC1936 study came from the Multi-Centre Research Ethics Committee for Scotland (MREC/01/0/56; 07/MRE00/58) and the Lothian Research Ethics Committee (LREC/2003/2/29). All participants, who were volunteers and received no financial or other reward, completed a written consent form before any testing or imaging took place.

##### 9.2 Image acquisition

Participants were imaged using a GE Signa Horizon HDxt 1.5 Tesla clinical scanner (General Electric, Milwaukee, WI, USA) using a self-shielding gradient set with maximum gradient strength of 33 mT/m and an 8-channel phased-array head coil. The full details of the imaging protocol can be found in (20). Briefly, the DTI examination consisted of 7 $T_2$-weighted (b = 0 s $mm^{-2}$) and sets of diffusion-weighted (b = 1000 s $mm^{-2}$) single-shot spin-echo echo-planar (EP) volumes acquired with diffusion gradients applied in 64 non-collinear directions (Jones et al., 2002). Volumes were acquired in the axial plane, with a field-of-view of 256 × 256 mm, contiguous slice locations, and image matrix and slice thickness designed to give 2 mm isotropic voxels. The repetition and echo time for each EP volume were 16.5 s and 98 ms respectively.

##### 9.3 DWI preprocessing

DTI data were converted from DICOM ((16)) to NIfTI-1 (http://nifti.-nimh.nih.gov/nifti-1) format using the TractoR package (http://www.tractor-mri.org.uk). FSL tools (http://www.fmrib.ox.ac.uk/fsl) were then used to extract the brain, remove bulk motion and eddy current induced distortions by registering all subsequent volumes to the first $T_2$-weighted EP volume (21), estimate the water diffusion tensor and calculate parametric maps of mean diffusivity (MD) and fractional anisotropy (FA) from its eigenvalues using DTIFIT.

#### 10 MAS

##### 10.1.1 Participants

Participants were drawn from Wave 2 of the Sydney Memory and Aging Study (MAS), a longitudinal study examining the predictors of cognitive decline in an elderly, non-demented, community-dwelling sample. They were recruited randomly through the electoral roll from two electorates of Eastern Sydney, Australia, where registration on the electoral roll is compulsory. Those who had a Wave 1 MRI scan were offered a 2year follow-up MRI at Wave 2. The study was approved by the Ethics Committee of the University of New South Wales and the details of the study have been published elsewhere (22).

##### 10.1.2 Image acquisition

All subjects were scanned using a Philips 3 Tesla Achieva Quasar Dual scanner (Philips Medical Systems) located at the Prince of Wales Medical Research Institute, Sydney. A single-shot echo-planar imaging (EPI) sequence (TR =7115 ms, TE = 70 ms) was used. Diffusion sensitizing gradients were applied along 32 non-collinear directions (b=1000 s/mm2), together with a non-diffusion-weighted acquisition (b = 0 s/mm2). For each DWI scan, 55 axial slices were collected. The field of view was 240mm × 240mm × 137.5mm with acquisition matrix 96 × 96 and zero filled into 240 × 240; and slice thickness 2.5 mm with no gap, yielding 1 mm ×1 mm × 2.5 voxels. Two extra non-diffusion-weighted (b = 0 s/mm2) EPIs were separately acquired and then combined with DTI scans for higher SNR.

##### 10.1.3 DWI preprocessing

DTI data were pre-processed with the FMRIB's Diffusion Toolbox (FDT) of the FMRIB's Software Library (FSL) version 5.0 (http://www.Fmrib.ox.ac.uk/fsl, Oxford Centre for Functional Magnetic Resonance Imaging of the Brain, Oxford University, UK). First, each subject's raw DTI data were corrected for head movement and eddy current distortions by linearly registering each diffusion-weighted volume to the reference T-weighted non-diffusion b0 image, using the FMRIB's linear image registration (FLIRT). A binary brain mask was created to remove the non-brain tissue using the Brain Extraction Tool (BET) in FSL. Then, diffusion tensor model was fitted to each imaging voxel of the pre-processed DTI data to derive Fractional anisotropy (FA) and diffusivity (AD, RD, MD) maps.

#### References

1. Smith SM, Jenkinson M, Woolrich MW, Beckmann CF, Behrens TEJ, Johansen-Berg H, Bannister PR, De Luca M, Drobnjak I, Flitney DE, et al. Advances in functional and structural MR image analysis and implementation as FSL. *NeuroImage* (2004) **23**:S208–S219. doi:10.1016/j.neuroimage.2004.07.051

2. Andersson JLR, Graham MS, Zsoldos E, Sotiropoulos SN. Incorporating outlier detection and replacement into a non-parametric framework for movement and distortion correction of diffusion MR images. *NeuroImage* (2016) **141**:556–572. doi:10.1016/j.neuroimage.2016.06.058

3. Andersson JLR, Sotiropoulos SN. An integrated approach to correction for off-resonance effects and subject movement in diffusion MR imaging. *NeuroImage* (2016) **125**:1063–1078. doi:10.1016/j.neuroimage.2015.10.019

4. Manjón JV, Coupé P, Martí-Bonmatí L, Collins DL, Robles M. Adaptive non-local means denoising of MR images with spatially varying noise levels: Spatially Adaptive Nonlocal Denoising. *J Magn Reson Imaging* (2010) **31**:192–203. doi:10.1002/jmri.22003

5. Mazoyer B, Mellet E, Perchey G, Zago L, Crivello F, Jobard G, Delcroix N, Vigneau M, Leroux G, Petit L, et al. BIL&GIN: A neuroimaging, cognitive, behavioral, and genetic database for the study of human brain lateralization. *NeuroImage* (2016) **124**:1225–1231. doi:10.1016/j.neuroimage.2015.02.071

6. Jenkinson M, Beckmann CF, Behrens TEJ, Woolrich MW, Smith SM. FSL. *NeuroImage* (2012) **62**:782–790. doi:10.1016/j.neuroimage.2011.09.015

7. Güllmar D, Haueisen J, Reichenbach JR. Analysis of *b* -value calculations in diffusion weighted and diffusion tensor imaging: *b* -Value Calculations. *Concepts Magn Reson Part A* (2005) **25A**:53–66. doi:10.1002/cmr.a.20031

8. Garyfallidis E, Brett M, Amirbekian B, Rokem A, van der Walt S, Descoteaux M, Nimmo-Smith I, Dipy Contributors. Dipy, a library for the analysis of diffusion MRI data. *Front Neuroinformatics* (2014) **8**: doi:10.3389/fninf.2014.00008

9. Paus T, Pausova Z, Abrahamowicz M, Gaudet D, Leonard G, Pike GB, Richer L. Saguenay Youth Study: A multi-generational approach to studying virtual trajectories of the brain and cardio-metabolic health. *Dev Cogn Neurosci* (2015) **11**:129–144. doi:10.1016/j.dcn.2014.10.003

10. Pausova Z, Paus T, Abrahamowicz M, Bernard M, Gaudet D, Leonard G, Peron M, Pike GB, Richer L, Séguin JR, et al. Cohort Profile: The Saguenay Youth Study (SYS). *Int J Epidemiol* (2017) **46**:1-14. doi:10.1093/ije/dyw023

11. Loeffler M, Engel C, Ahnert P, Alfermann D, Arelin K, Baber R, Beutner F, Binder H, Brähler E, Burkhardt R, et al. The LIFE-Adult-Study: objectives and design of a population-based cohort study with 10,000 deeply phenotyped adults in Germany. *BMC Public Health* (2015) **15**:691. doi:10.1186/s12889-015-1983-z

12. Smith SM. Fast robust automated brain extraction. *Hum Brain Mapp* (2002) **17**:143–155. doi:10.1002/hbm.10062

13. Lohmann G, MuÈller K, Bosch V, Mentzel H, Hessler S, Chen L, Zysset S. LipsiaÐa new software system for the evaluation of functional magnetic resonance images of the human brain. *Comput Med Imaging Graph* (2001)9.

14. Caspers S, Moebus S, Lux S, Pundt N, Schütz H, Mühleisen TW, Gras V, Eickhoff SB, Romanzetti S, Stöcker T, et al. Studying variability in human brain aging in a population-based German cohortâ€"rationale and design of 1000BRAINS. *Front Aging Neurosci* (2014) **6**: doi:10.3389/fnagi.2014.00149

15. Koster J, Rahmann S. Snakemake--a scalable bioinformatics workflow engine. *Bioinformatics* (2012) **28**:2520–2522. doi:10.1093/bioinformatics/bts480

16. Schmidt R, Lechner H, Fazekas F, Niederkorn K, Reinhart B, Grieshofer P, Horner S, Offenbacher H, Koch M, Eber B, et al. Assessment of cerebrovascular risk profiles in healthy-persons - definition of research goals and the Austrian Stroke Prevention Study (ASPS). *Neuroepidemiology* (1994) **13**:308–313. doi:10.1159/000110396

17. Schmidt R, Fazekas F, Kapeller P, Schmidt H, Hartung HP. MRI white matter hyperintensities - Three-year follow-up of the Austrian stroke prevention study. *Neurology* (1999) **53**:132–139. doi:10.1212/WNL.53.1.132

18. Sachdev PS, Lammel A, Trollor JN, Lee T, Wright MJ, Ames D, Wen W, Martin NG, Brodaty H, Schofield PR, et al. A comprehensive neuropsychiatric study of elderly twins: The Older Australian Twins Study. *Twin Res Hum Genet* (2009) **12**:573–582. doi:10.1375/twin.12.6.573

19. Taylor AM, Pattie A, Deary IJ. Cohort Profile Update: The Lothian Birth Cohorts of 1921 and 1936. *Int J Epidemiol* (2018) **47**:1042–1042r. doi:10.1093/ije/dyy022

20. Wardlaw JM, Bastin ME, Valdés Hernández MC, Maniega SM, Royle NA, Morris Z, Clayden JD, Sandeman EM, Eadie E, Murray C, et al. Brain aging, cognition in youth and old age and vascular disease in the Lothian Birth Cohort 1936: Rationale, Design and Methodology of the Imaging Protocol. *Int J Stroke* (2011) **6**:547–559. doi:10.1111/j.1747-4949.2011.00683.x

21. Jenkinson M, Smith S. A global optimisation method for robust affine registration of brain images. *Med Image Anal* (2001) **5**:143–156. doi:10.1016/S1361-8415(01)00036-6

22. Sachdev PS, Brodaty H, Reppermund S, Kochan NA, Trollor JN, Draper B, Slavin MJ, Crawford J, Kang K, Broe GA, et al. The Sydney Memory and Ageing Study (MAS): methodology and baseline medical and neuropsychiatric characteristics of an elderly

epidemiological non-demented cohort of Australians aged 70–90 years. *Int Psychogeriatr* (2010) **22**:1248–1264. doi:10.1017/S1041610210001067
